# SARS-CoV-2 Nonstructural Proteins 3 and 4 tune the Unfolded Protein Response

**DOI:** 10.1101/2023.04.22.537917

**Authors:** Jonathan P. Davies, Athira Sivadas, Katherine R. Keller, Richard J.H. Wojcikiewicz, Lars Plate

**Affiliations:** Department of Biological Sciences, Vanderbilt University, Nashville, TN; Department of Pharmacology, SUNY Upstate Medical University, Syracuse, NY; Department of Chemistry, Vanderbilt University, Nashville, TN; Department of Pathology, Microbiology and Immunology, Vanderbilt University Medical Center, Nashville, TN

**Keywords:** coronavirus, proteomics, stress response, nonstructural protein, ATF6, PERK

## Abstract

Coronaviruses (CoV), including SARS-CoV-2, modulate host proteostasis through activation of stress-responsive signaling pathways such as the Unfolded Protein Response (UPR), which remedies misfolded protein accumulation by attenuating translation and increasing protein folding capacity. While CoV nonstructural proteins (nsps) are essential for infection, little is known about the role of nsps in modulating the UPR. We characterized the impact of SARS-CoV-2 nsp4, a key driver of replication, on the UPR using quantitative proteomics to sensitively detect pathway-wide upregulation of effector proteins. We find nsp4 preferentially activates the ATF6 and PERK branches of the UPR. Previously, we found an N-terminal truncation of nsp3 (nsp3.1) can suppress pharmacological ATF6 activation. To determine how nsp3.1 and nsp4 tune the UPR, their co-expression demonstrated that nsp3.1 suppresses nsp4-mediated PERK, but not ATF6 activation. Re-analysis of SARS-CoV-2 infection proteomics data revealed time-dependent activation of PERK targets early in infection, which subsequently fades. This temporal regulation suggests a role for nsp3 and nsp4 in tuning the PERK pathway to attenuate host translation beneficial for viral replication while avoiding later apoptotic signaling caused by chronic activation. This work furthers our understanding of CoV-host proteostasis interactions and highlights the power of proteomic methods for systems-level analysis of the UPR.

## Introduction

The coronavirus SARS-CoV-2 has caused the COVID-19 pandemic, resulting in more than 6.8 million deaths to date^1^. Coronaviruses (CoV) require host cell translation machinery and endoplasmic reticulum (ER) derived membranes for replication. CoV infection is known to induce ER stress and activate the Unfolded Protein Response (UPR)^2–4^. The UPR alleviates stress from accumulation of misfolded proteins in the ER lumen by temporarily suppressing global protein translation, increasing the production of ER chaperones, expanding ER membrane synthesis, and if stress persists, triggering apoptosis. The UPR is composed of three branches that signal downstream of their respective ER-membrane localized stress sensors: 1. protein kinase R-like ER kinase (PERK), 2. Inositol requiring enzyme 1α (IRE1α), and 3. activating transcription factor 6 (ATF6) pathways ^5,6^.

Activation of PERK leads to the phosphorylation of eukaryotic initiation factor 2α (eIF2α) and attenuation of global protein translation to prevent further accumulation of misfolded proteins in the ER. A select group of proteins are translated under these conditions, including activating transcription factor 4 (ATF4), which upregulates expression of various genes involved in protein folding, antioxidant response, and the pro-apoptotic transcription factor C/EBP Homologous Protein (CHOP)^7,8^. IRE1α has endoribonuclease activity which, upon ER stress, cleaves X-box-binding protein 1 (*XBP1*) transcripts, leading to splicing and translation of the XBP1s transcription factor and increased gene expression of ER protein chaperones, translocation and secretion factors, and components of ER-associated degradation (ERAD), as well as ER biogenesis^9^. Lastly, ATF6 is translocated to the Golgi upon ER stress, where it is cleaved by site-1 and site-2-proteases. The N-terminal fragment (ATF6p50) is a transcription factor which initiates upregulation of various protein folding, secretion, and degradation factors, as well as expansion of the ER^5,6,10,11^.

Coronavirus replication requires extensive production, folding, and modification of viral proteins^12^, as well as alteration of ER-derived membranes to form double-membrane vesicles (DMVs) for replication sites. These processes can trigger ER stress and activate the UPR^13^. Indeed, a previous study found that all three branches of the UPR are activated during SARS-CoV-2 infection and that expression of SARS-CoV-2 Spike or orf8 protein is sufficient to activate the UPR^3^. SARS-CoV proteins orf3a and orf8ab were also found to activate the PERK and ATF6 pathways respectively^14,15^. A related betacoronavirus, mouse hepatitis virus (MHV), triggers activation of XBP1 and PERK pathways while also hindering production of certain UPR-responsive genes such as CHOP^4^. This previous work shows that coronaviruses can modulate stress responses at multiple phases and emphasizes the need for downstream proteome measurements to elucidate the consequences in the host cell.

Surprisingly, relatively little is known about the impact of CoV nonstructural proteins (nsps) on the UPR. Nsps are the first viral proteins translated during infection, rewire the host cell, and replicate the viral genome. In particular, three transmembrane nsps (nsp3, nsp4, and nsp6) are responsible for DMV formation from ER-derived membranes^16^. Nsp3 contains a papain-like protease domain (PL2^pro^), which cleaves the orf1a/b polypeptide and also possesses deubiquination/de-ISGylation activity^17,18^. Nsp4 is a glycoprotein containing four transmembrane domains and plays a key role in membrane reorganization^19,20^.

We have previously characterized the interactomes of nsp3 and nsp4 CoV homologs and found that both proteins have evolutionary conserved interactions with several ER proteostasis factors^21,22^. We identified an interaction between an N-terminal fragment of SARS-CoV-2 nsp3 (nsp3.1) and ATF6 and showed that nsp3.1 can suppress pharmacologic activation of the ATF6 pathway^22^. Given the known role of nsp4 in host membrane alteration, we sought to elucidate if nsp4 activates or suppresses the UPR and whether nsp4 may act in concert with nsp3 to tune UPR activation. In particular, we leveraged a quantitative proteomics approach to characterize the upregulation of genes known to be transcriptionally activated by the UPR^6,23,24^. Importantly, this approach enables sensitive measurement of pathway-wide changes in effector proteins and accounts for viral protein-mediated regulation that may occur downstream of transcriptional activation. A more precise understanding of the role of CoV nsps in modulating the UPR will further our knowledge of how coronavirus replication manipulates host proteostasis pathways during infection.

### Experimental Section

#### DNA Constructs

SARS-CoV-2 nsp4-FT, nsp3-FT, FT-nsp2, and orf8-FT (Wuhan-Hu-1 MN908947) were codon-optimized and cloned into a pcDNA-(+)-C-DYK vector (Genscript) or pcDNA-(+)-N-DYK vector for nsp2 (Genscript). An N-terminal truncation nsp3.1-FT (residues 1-749) was generated as previously described ^21^. Nsp4-ST and orf8-ST constructs were generated through the NEB HiFi Assembly system. In brief, the FLAG-tag from nsp4-FT was removed through amplification with primers 3 & 4 respectively. A 2xStrepTag was amplified from a SARS-CoV-2 nsp2-ST in a pLVX-EF1alpha plasmid construct (kind gift from Dr. Nevan Krogan, University of California, San Francisco) using primers 1 & 2. The linear nsp4 product was then combined with the 2xStrepTag fragment via HiFi assembly (1:2 vector to insert ratio). The orf8-ST construct was made by amplifying out the orf8 gene from the pcDNA-(+)-C-DYK vector using primers 5 & 6. A pLVX-EF1alpha-SARS-CoV-2-nsp2-2xStrepTag plasmid was linearized using primers 7 & 8, retaining the vector backbone and 2xStrepTag while removing the nsp2 gene. These were then combined via HiFi assembly (1:2 vector to insert ratio). Plasmids were verified by sequencing (Genewiz).

#### Primers

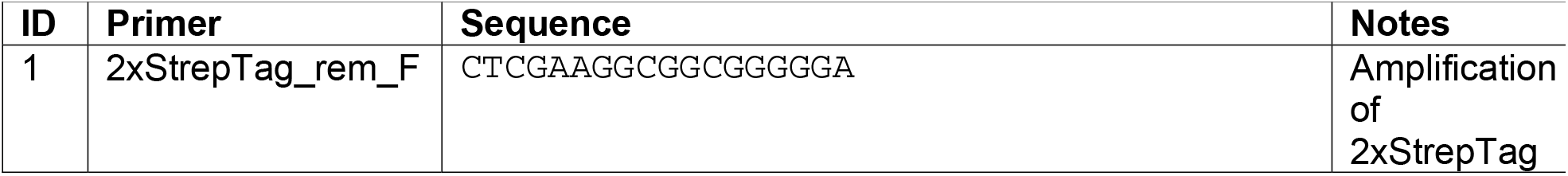
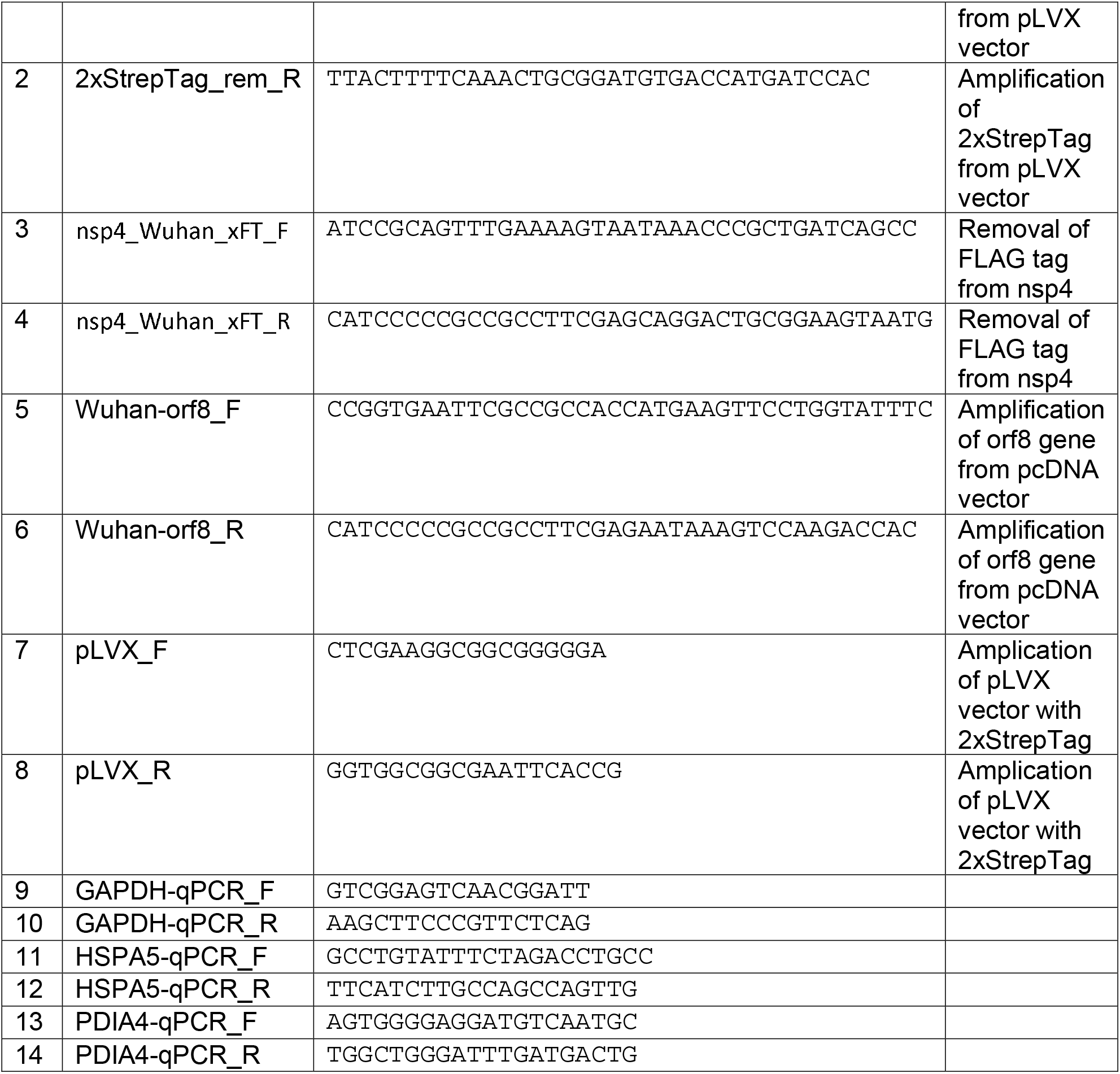

#### Cell Culture and Transfection

HEK293T cells were generously provided by Dr. Joseph Genereux (University of California, Riverside). A549 lung epithelial cells were obtained from ATCC (CCL-185). Cell lines were tested regularly for mycoplasma. HEK293T and A549 cells were maintained at 37°C, 5% CO2 in high glucose Dulbecco’s Modified Eagle Growth Medium with 10% fetal bovine serum, 1% glutamine, and 1% penicillin/streptomycin (DMEM-10). Cells were seeded into 6-well plates at a concentration of 4 × 10^5^ cells/well. HEK293T cells were transfected dropwise, twenty-four hours after seeding, using a calcium phosphate cocktail (1.5 µg total DNA, 0.25 M CaCl_2_, 1x HBS (137 mM NaCl, 5 mM KCl, 0.7 mM Na_2_HPO_4,_ 7.5 mM D-glucose, 21 mM HEPES)). The media was changed 18 hours later with fresh DMEM-10 media. A549 cells were seeded in antibiotic-free DMEM-10 and 24 h later were transfected with FuGENE 4K transfection reagent (FuGENE #E5911) (1.5 µg DNA with 4.5 µL FuGENE 4K) in plain Optimem media. Samples were treated with DMSO or Tunicamycin (Tm, 1 µg/mL) for 16 h (protein analysis) or 6 h (RNA analysis) prior to harvest in described experiments. GFP controls were used to assess transfection efficiency, which averaged 70-80% in HEK293T cells and 40-50% in A549 cells. Transfected cells were harvested by cell scraping in cold 1 mM EDTA in PBS over ice, 40 h post-transfection to enable robust expression of viral proteins. To measure CHOP levels, HEK293T cells were transfected twenty-four hours after seeding, using 1µg cDNAs and 6 µl of 1mg/ml PEI (pre-mixed in 50 µl of serum-free culture medium). Samples were treated with Tunicamycin (Tm, 5 µg/mL) for 4h or 20h prior to harvest.

#### RT-qPCR

HEK293T cells were transfected and harvested as described previously. Cellular RNA was extracted using the Zymo QuickRNA miniprep kit (#R1055) and 500 ng total cellular RNA was reverse transcribed into cDNA using random hexamer primers (IDT #51-01-18-25), oligo-dT 15mer primers (IDT # 51-01-15-05), and Promega M-MLV reverse transcriptase (#M1701). qPCR analysis was carried out using BioRad iTaq Universal SYBR Green Supermix (#1725120), added to respective primers (primers 9-14) for target genes and reactions were run in 96-well plates on a BioRad CFX qPCR instrument. Conditions used for amplification were 95°C, 2 min, 45 repeats of 95°C, 10 s and 60°C, 30 s. A melting curve was generated in 0.5°C intervals from 65 to 95°C. Cq values were calculated by the BioRad CFX Maestro software and transcripts were normalized to a housekeeping gene (*GAPDH*). All measurements were performed in technical duplicate; technical duplicates were averaged to form a single biological replicate.

#### Western Blot Analysis

HEK293T cells were lysed on ice in TNI buffer (50mM Tris pH 7.5, 150mM NaCl, 0.5% IGEPAL-CA-630) with Roche c0mplete EDTA-free protease inhibitor (#4693132001) for 15 minutes and then sonicated for 10 minutes in a water bath at room temperature. A549 cells were lysed on ice in RIPA buffer (50 mM Tris pH 7.5, 150 mM NaCl, 0.1% SDS, 1% Triton X-100, 0.5% deoxycholate) with Roche c0mplete EDTA-free protease inhibitor (#4693132001) for 15 minutes. Lysates were spun down at 21.1k xg for 15 minutes at 4°C. Samples were added to 6x Laemelli buffer (12% SDS, 125 mM Tris, pH 6.8, 20% glycerol, bromophenol blue, 100 mM DTT) and heated at 37°C for thirty minutes. The samples were then run on an SDS-PAGE gel and transferred to a PVDF membrane for Western blotting. M2 anti-FLAG (Sigma Aldrich, F1804), anti-KDEL (Enzo ADI-SPA-827-F), anti-PDIA4 (ProteinTech 14712-1-AP), THE anti-Strep II tag FITC (Genescript, A01736-100), and anti-GAPDH (GeneTex, GTX627408) antibodies were used to probe Western blots at a 1:1000 dilution in TBS blocking buffer (0.1% Tween, 5% BSA, 0.1% Sodium Azide).

To measure CHOP protein levels, HEK293T cells were lysed on ice in 1% CHAPS lysis buffer (50 mM Tris H-Cl, 150 mM NaCl, 1mM EDTA, 1% CHAPS, 10 µM pepstatin, 0.2 mM phenylmethylsulfonyl fluoride, 0.2 µM soybean trypsin inhibitor, and 1mM DTT; pH 8.0) for 30 minutes. Lysates were spun down at 16,000 xg for 10 minutes at 4°C. Samples were added to 4x gel-loading buffer (50mM Tris/HCl (pH 6.8), 100mM DTT, 2% SDS, 0.1% Bromophenol Blue, 10% glycerol, 100 mM DTT) and heated at 37°C for thirty minutes. The samples were then run on an 13% SDS-PAGE gel and transferred to a nitrocellulose membrane for Western blotting. M2 anti-FLAG (Sigma Aldrich, F1804), anti-BiP #3177 (Cell Signaling Technology), anti-CHOP #7351, and anti-GAPDH #365062 (Santa Cruz Biotechnology Inc.), antibodies were used to probe Western blots in TBS blocking buffer (0.1% Tween, 4% BSA, 0.1% Sodium Azide). GAPDH signal was used to normalize band intensities. Protein expression on Western blots was quantified using ImageLab.

#### Proteomics Experimental Design

HEK293T UPR analysis combined 2-3 biological replicates into 1 individual MS run (3x GFP, 3x nsp3.1-FT+GFP, 3x nsp4-ST+GFP, 3x nsp3.1-FT+nsp4-ST, 2x orf8-ST+GFP, 2x orf8-ST+nsp3.1-FT). A549 UPR analysis combined 4 biological replicates into 1 individual MS run (4x GFP, 4x GFP+Tm, 4x nsp4-FT).

#### Sample Preparation for Mass Spectrometry

Samples were harvested and lysed as described in *Western Blot Analysis*. Protein concentration was quantified using 1x BioRad Protein Assay Dye (#5000006) and 20 µg of protein from each sample was prepared for mass spectrometry. Proteins were precipitated via mass spectrometry grade methanol:chloroform:water (in a 3:1:3 ratio) and washed three times with methanol. Each wash was followed by a 2-minute spin at 10,000xg at room temperature. Protein pellets were air dried 30-45 min and resuspended in 5 µL of 1% Rapigest SF (Waters #186002122). Resuspended proteins were diluted with 32.5 µL water and 10 µL 0.5 M HEPES (pH 8.0), then reduced with 0.5 µL of freshly made 0.5 M TCEP for 30 minutes at room temperature. Samples were then alkylated with 1 µL of fresh 0.5 M iodoacetamide (freshly made) for 30 minutes at room temperature in the dark and digested with 0.5 µg Pierce Trypsin/Lys-C (Thermo Fisher # A40007) overnight at 37°C shaking. Peptides were diluted to 60 µL with LC/MS-grade water and labeled using TMTpro labels (Thermo Scientific # A44520) for 1 h at room temperature. Labeling was quenched with the addition of fresh ammonium bicarbonate (0.4% v/v final) for 1 h at room temperature. Samples were then pooled, acidified to pH < 2.0 using formic acid, concentrated to 1/6^th^ original volume via Speed-vac, and diluted back to the original volume with buffer A (95% water, 5% acetonitrile, 0.1% formic acid). Cleaved Rapigest products were removed by centrifugation at 17,000xg for 30 minutes and supernatant transferred to fresh tubes.

#### MudPIT LC-MS/MS Analysis

Alternating layers of 1.5cm Aqua 5 µm C18 resin (Phenomenex # 04A-4299), 1.5cm Luna 5 µm SCX resin (Phenomenex # 04A-4398), and 1.5cm Aqua 5 µm C18 resin were packed to make triphasic MudPIT columns as described previously^25^. TMT-labeled samples (20 µg) were loaded onto the microcapillaries via a high-pressure chamber, followed by a 30-minute wash in buffer A (95% water, 5% acetonitrile, and 0.1% formic acid). The MudPIT columns were installed on the LC column switching valve and followed by a 20cm fused silica microcapillary column filled with Aqua C18, 3µm resin (Phenomenex # 04A-4311) ending in a laser-pulled tip. Columns were washed in the same way as the MudPIT capillaries prior to use. Liquid chromatography (Ultimate 3000 nanoLC system) was used to fractionate the peptides online and then analyzed via an Exploris480 mass spectrometer (Thermo Fisher). MudPIT runs were carried out by 10µL sequential injections of 0, 10, 20, 30, 40, 50, 60, 70, 80, 90, 100% buffer C (500mM ammonium acetate, 94.9% water, 5% acetonitrile, 0.1% formic acid), followed by a final injection of 90% C, 10% buffer B (99.9% acetonitrile, 0.1% formic acid v/v). Each injection was followed by a 130 min gradient using a flow rate of 500nL/min (0-6 min: 2% buffer B, 8 min: 5% B, 100 min: 35% B, 105min: 65% B, 106-113 min: 85% B, 113-130 min: 2% B). ESI was performed directly from the tip of the microcapillary column using a spray voltage of 2.2 kV, an ion transfer tube temperature of 275°C and an RF Lens of 40%. MS1 spectra were collected using a scan range of 400-1600 m/z, 120k resolution, AGC target of 300%, and automatic injection times. Data-dependent MS2 spectra were obtained using a monoisotopic peak selection mode: peptide, including charge state 2-7, TopSpeed method (3s cycle time), isolation window 0.4 m/z, HCD fragmentation using a normalized collision energy of 36%, 45k resolution, AGC target of 200%, automatic maximum injection times, and a dynamic exclusion (20 ppm window) set to 60s.

#### Peptide identification and quantification

Identification and quantification of peptides were performed in Proteome Discoverer 2.4 (Thermo Fisher) using the SwissProt human database (TaxID 9606, released 11/23/2019; 42,252 entries searched) with nsp3.1, nsp4, and orf8 fragment sequences (3 entries) manually added (42,255 total entries searched). Searches were conducted with Sequest HT using the following parameters: trypsin cleavage (maximum two missed cleavages), minimum peptide length 6 AAs, precursor mass tolerance 20 ppm, fragment mass tolerance 0.02 Da, dynamic modifications of Met oxidation (+15.995 Da), protein N-terminal Met loss (−131.040 Da), and protein N-terminal acetylation (+42.011 Da), static modifications of TMTpro (+304.207 Da) at Lys, and N-termini and Cys carbamidomethylation (+57.021 Da). Peptide IDs were filtered using Percolator with an FDR target of 0.01. Proteins were filtered based on a 0.01 FDR, and protein groups were created according to a strict parsimony principle. TMT reporter ions were quantified considering unique and razor peptides, excluding peptides with co-isolation interference greater than 25%. Peptide abundances were normalized based on total peptide amounts in each channel, assuming similar levels of background. Protein quantification used all quantified peptides. Post-search filtering was carried out to include only proteins with two or more identified peptides.

## Data Availability

The mass spectrometry proteomics data have been deposited to the ProteomeXchange Consortium via the PRIDE partner repository with the data set identifier PXD039797. All other necessary data are contained within the manuscript.

*The reviewers can access the data using the account information below:*

*Username:*

*Password: m1fxQW9n*

## Results and Discussion

### SARS-CoV-2 nsp4 upregulates expression of UPR reporter proteins

Given the role of nsp4 in modulating host ER membranes during infection, we hypothesized that nsp4 may induce ER stress and subsequently activate the UPR pathway. To test this, we exogenously expressed a C-terminally FLAG-tagged SARS-CoV-2 nsp4 construct (nsp4-FT, **Fig. 1a**) in HEK293T cells as previously reported^21^ and measured transcript and protein expression of several UPR branch markers. Tdtomato (Tdt) expression was used as a negative control to account for any transfection-induced cell stress and benchmark the basal state of UPR marker levels. Importantly, previous work has shown transient expression of fluorescent, cytosolic proteins does not induce ER stress and the UPR^26^. Treatment with tunicamycin (Tm, 1 µg/mL, 6 h) was used as a positive control for strong, general UPR activation. Samples were harvested 40 hours post-transfection, to ensure robust expression of viral proteins. Using RT-qPCR, we found that nsp4-FT expression led to a moderate but signification upregulation of *HSPA5* transcripts and upregulation, though not significantly, of *PDIA4* transcripts, both markers for ATF6 branch-activation ^6,11,27^ (**Fig. 1b**).

**Figure 1.**
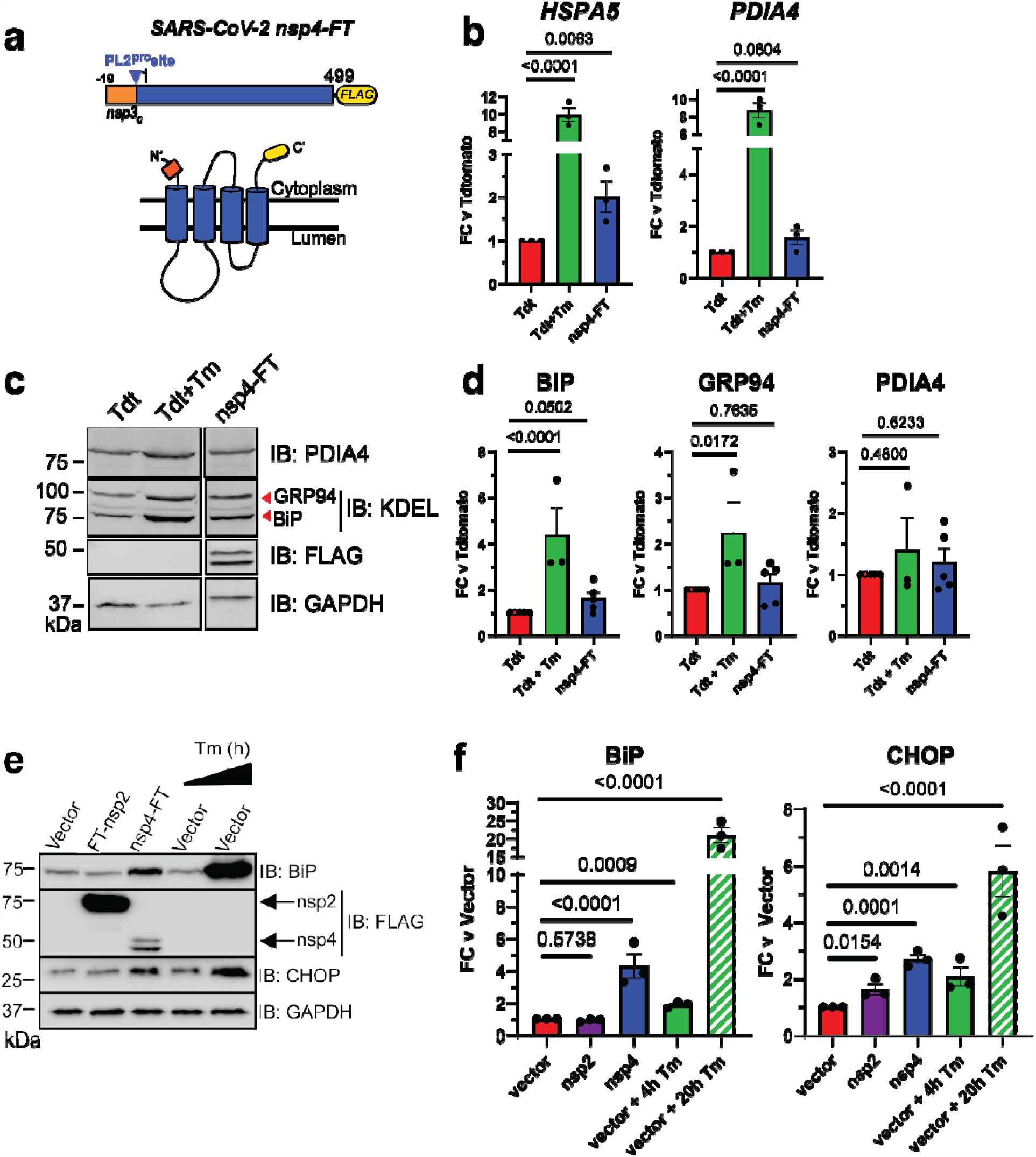
SARS-CoV-2 nsp4 upregulates expression of UPR reporter proteins. **a)** SARS-CoV-2 nsp4 expression construct and membrane topology, containing a C-terminal FLAG-tag (nsp4-FT) and an N-terminal segment of the last 19 C-terminal amino acids in nsp3 for optimal membrane insertion^21^. Construct contains the native PL2^pro^ protease cleavage site. **b)** RT-qPCR of ATF6 pathway activation reporters *HSPA5* and *PDIA4* in the presence of SARS-CoV-2 nsp4-FT, normalized to *GAPDH* transcripts and compared to basal levels (GFP). Treatment with 1 µg/mL Tunacimycin (Tm) for 6 h was used as a positive control. n = 3, mean ±SEM, One-way ANOVA with Benjamini, Krieger and Yekutieli multiple testing correction, p<0.05 considered statistically significant. **c)** Representative Western blot of ATF6 protein markers (PDIA4, GRP94, BiP) in the presence of Tdtomato (Tdt), SARS-CoV-2 nsp4-FT, or treatment with 1 µg/mL Tm (16 h), with GAPDH as a housekeeping gene for loading control. Displayed blot sections are from the same blot image and exposure settings (see **Supplemental Fig. S1** for all quantified, original blots). **d)** Quantification of Tdtomato (Tdt), and nsp4-FT in Western blots in **(c)**, normalized to GAPDH band intensities, and compared to basal levels (Tdt). n = 3-5, mean ±SEM, One-way ANOVA with Benjamini, Krieger and Yekutieli multiple testing correction, p<0.05 considered statistically significant. **e)** Representative Western blot of CHOP and BiP in the presence of SARS-CoV-2 FT-nsp2, nsp4-FT, or treatment with 5 µg/mL Tm for short (4h) or long (20h) time points. See **Supplemental Fig. S2** for all quantified, original blots. **f)** Quantification of Western blots in **(e)**, normalized to GAPDH band intensities and compared to basal control (vector). n = 3, mean ±SEM. One-way ANOVA with Benjamini, Krieger and Yekutieli multiple testing correction, p<0.05 considered statistically significant.

To probe downstream protein expression of ATF6 UPR markers, we measured BiP, GRP94, and PDIA4 levels by Western blot in the presence of nsp4-FT (**Fig. 1c,d,e,f, Supplemental Fig. S1, S2**). These proteins have been extensively used in prior studies as Western blot markers for ATF6^28^. Quantification of the control and nsp4-FT lanes show that nsp4 induces a modest upregulation of ATF6 reporter proteins (**Fig. 1d,f**). We also found that nsp4-FT induces strong upregulation of a PERK pathway marker, CHOP^29^ (**Fig. 1e,f, Supplemental Fig. S2**), while expression of SARS-CoV-2 FT-nsp2 (a cytosolic viral protein) does not. These results indicate that SARS-CoV-2 nsp4 activates the UPR, albeit at milder levels than the toxic ER stressor tunicamycin, but still at biologically-relevant levels.

### Nsp4 upregulates the ATF6 and PERK pathways as measured by quantitative proteomics

While measuring a select few protein markers by Western blot provides some indication of UPR activation, a more precise and pathway-wide approach is required to comprehensively characterize downstream modulation of the cellular proteome. To this end, we analyzed the global proteome of HEK293T cells transfected with SARS-CoV-2 nsp4-StrepTag (nsp4-ST) using tandem mass spectrometry (LC-MS/MS) with TMTpro isobaric tags to quantify protein abundance (**Fig. 2a, Supplemental Tables S1, S2, S3**). Transfection of GFP was used as a control to indicate the basal state of the UPR (-nsp4). Samples were normalized based on global peptide abundance (**Supplemental Fig. S3**). To measure pathway-wide changes of each UPR branch, we used a previously defined set of genes that have been shown to be transcriptionally upregulated upon stress-independent activation by RNA-Seq analysis^6,24,30^ (**Supplemental Table S1)**. By quantifying proteome changes in these pathway reporters, we can account for viral protein-mediated alterations in translation or degradation of proteostasis factors that may occur downstream of transcriptional activation.

**Figure 2.**
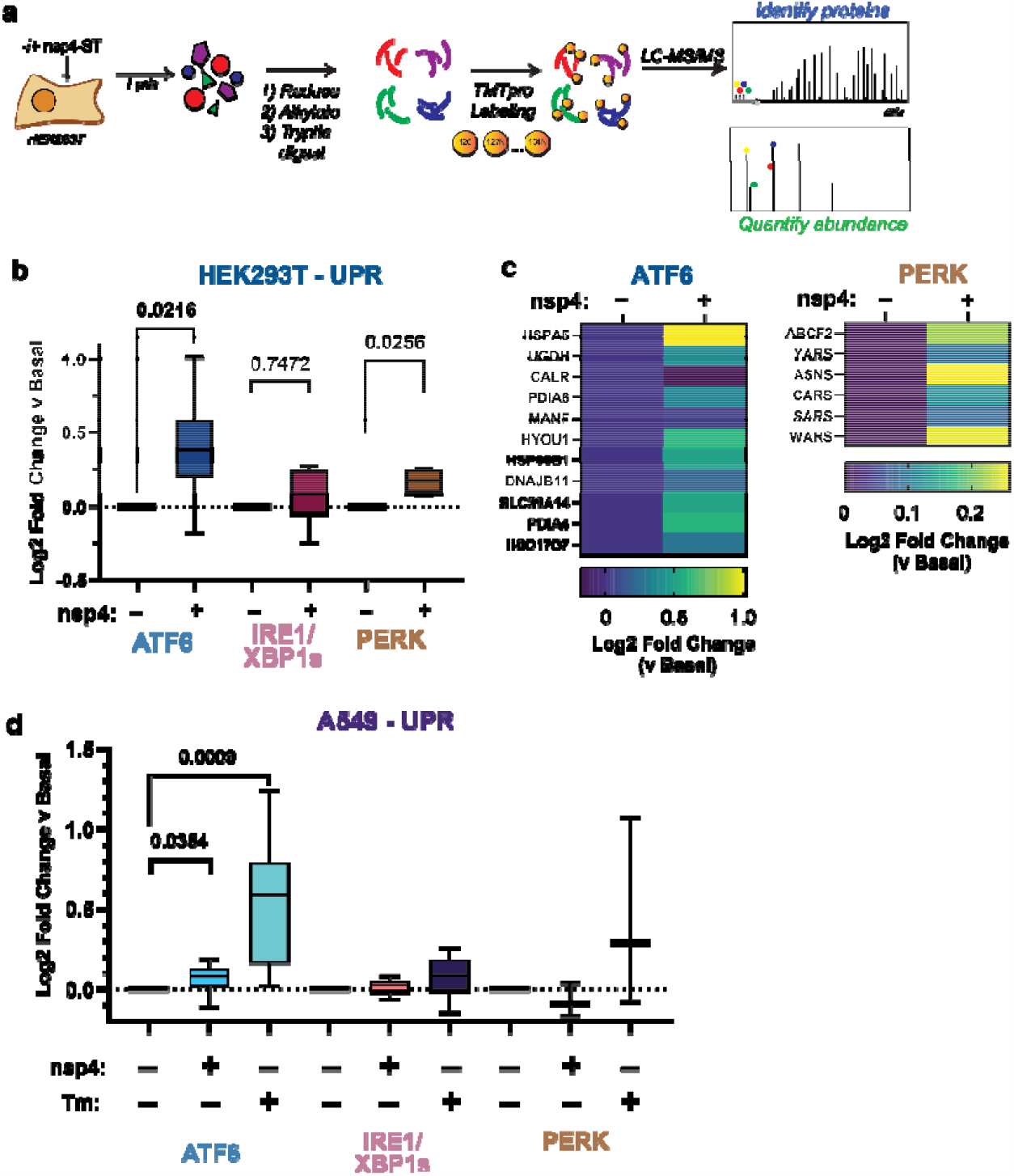
SARS-CoV-2 nsp4 upregulates the ATF6 and PERK pathways as measured by quantitative proteomics. **a)** Experimental schematic to measure UPR induction by SARS-CoV-2 nsp4-ST expression in HEK293T cells using tandem mass spectrometry (LC/MS-MS) and TMTpro-based quantification. **b)** Log2 fold change of UPR branch protein markers in the absence (GFP) or presence of SARS-CoV-2 nsp4-ST, as outlined in **(a)** assessed by TMT reporter ion intensities. Values represent individual protein markers previously defined by RNA-seq analysis to be transcriptionally upregulated upon stress-independent activation of respective UPR branches ^6,24,30^. Box-and-whisker plot shows median, 25^th^ and 75^th^ quartiles, and minimum and maximum values. n = 3 biological replicates, 1 MS run, one-way ANOVA with Geisser–Greenhouse correction and post-hoc Tukey’s multiple comparison test was used to test significance, *p*<0.05 considered significant. See **Supplemental Tables S2, S3** for mass spectrometry data set. **c)** Heatmap of individual log2 fold change for ATF6 and PERK protein markers in the absence or presence of SARS-CoV-2 nsp4-ST. **d)** Log2 fold change of UPR branch protein markers in A549 lung epithelial cells the presence of SARS-CoV-2 nsp4-FT or Tunicamycin (1 µg/mL, 16 h) compared to basal control (GFP). UPR branch protein markers were defined as in **(b)**. Box-and-whisker plot shows median, 25^th^ and 75^th^ quartiles, and minimum and maximum values. n = 4 biological replicates, 1 MS run, one-way ANOVA with Geisser–Greenhouse correction and post-hoc Tukey’s multiple comparison test was used to test significance, *p*<0.05 considered significant. See **Supplemental Tables S4, S5** for mass spectrometry data set.

We compared UPR pathway upregulation in the absence or presence of nsp4 by examining the distribution of target upregulation for each pathway (**Fig. 2b**). We found that both the ATF6 and PERK pathways were significantly upregulated.

We also examined the abundances of individual proteins identified within the ATF6 and PERK pathways (**Fig. 2c**). There was a heterogeneous ATF6 response to nsp4 expression, with some proteins (DNAJB11 and MANF) displaying relatively small change while others (HSPA5, PDIA4, HYOU1, HSP90B1) showed much higher upregulation. Comparing to previously published proteomics dataset^22^, we find nsp4-ST induces higher expression of ATF6 markers over basal levels compared to a specific ATF6 pharmacological activator, compound **147**, but to a lesser extent than Tm treatment (**Supplemental Fig. S4**). We found that the PERK pathway upregulation by nsp4 was largely dominated by changes in ASNS and WARS abundance (**Fig. 2c**).

To determine if this phenotype extends to disease-relevant cell models, we expressed SARS-CoV-2 nsp4-FT in A549 lung epithelial cells and measured changes in UPR markers via TMTpro LC/MS-MS (**Fig. 2d, Supplemental Fig. S3, Supplemental Tables S4, S5**). As a positive control for UPR activation, we included samples treated with Tm (1 µg/mL, 16 h). ATF6 protein markers are moderately but statistically significantly increased in the presence of nsp4, though to a lesser extent than in HEK293T cells, indicative of some cell-type specific effects. Together, these results indicate SARS-CoV-2 nsp4 activates an ATF6 and PERK response characterized by substantial ER chaperone upregulation, though to a lesser extent compared to the potent ER stressor, tunicamycin.

### SARS-CoV-2 nsp3.1 suppresses nsp4-induced PERK activation, but not ATF6 activation

Nsp4 and nsp3 are key inducers of double-membrane vesicle (DMV) formation during CoV replication^16,19^, which, along with viral protein production, may induce ER stress and the UPR. We previously showed an N-terminal fragment of nsp3 (nsp3.1) suppresses pharmacological ATF6 activation^22^. Therefore, we sought to test whether nsp3.1 might dampen nsp4-induced UPR activation, providing a form of viral tuning of the host UPR. To this end, we co-transfected HEK293T cells with SARS-CoV-2 nsp4-ST and nsp3.1-FT plasmids in equal amounts (0.75 µg) and measured the effect on ATF6 protein reporters by Western blot and UPR markers by mass spectrometry (**Fig. 3a, Supplemental Tables S2, S3**). In addition, we tested if nsp3.1 could suppress UPR activation induced by SARS-CoV-2 orf8-ST, another viral protein known to activate the UPR^3^. A GFP transfection served as a control for basal UPR state. To ensure that all samples were expressing similar amounts of protein, individual viral protein transfections were supplemented with equal amounts of GFP-expression plasmid (0.75 µg each). All transfections used 1.5 µg DNA plasmid in total.

**Figure 3.**
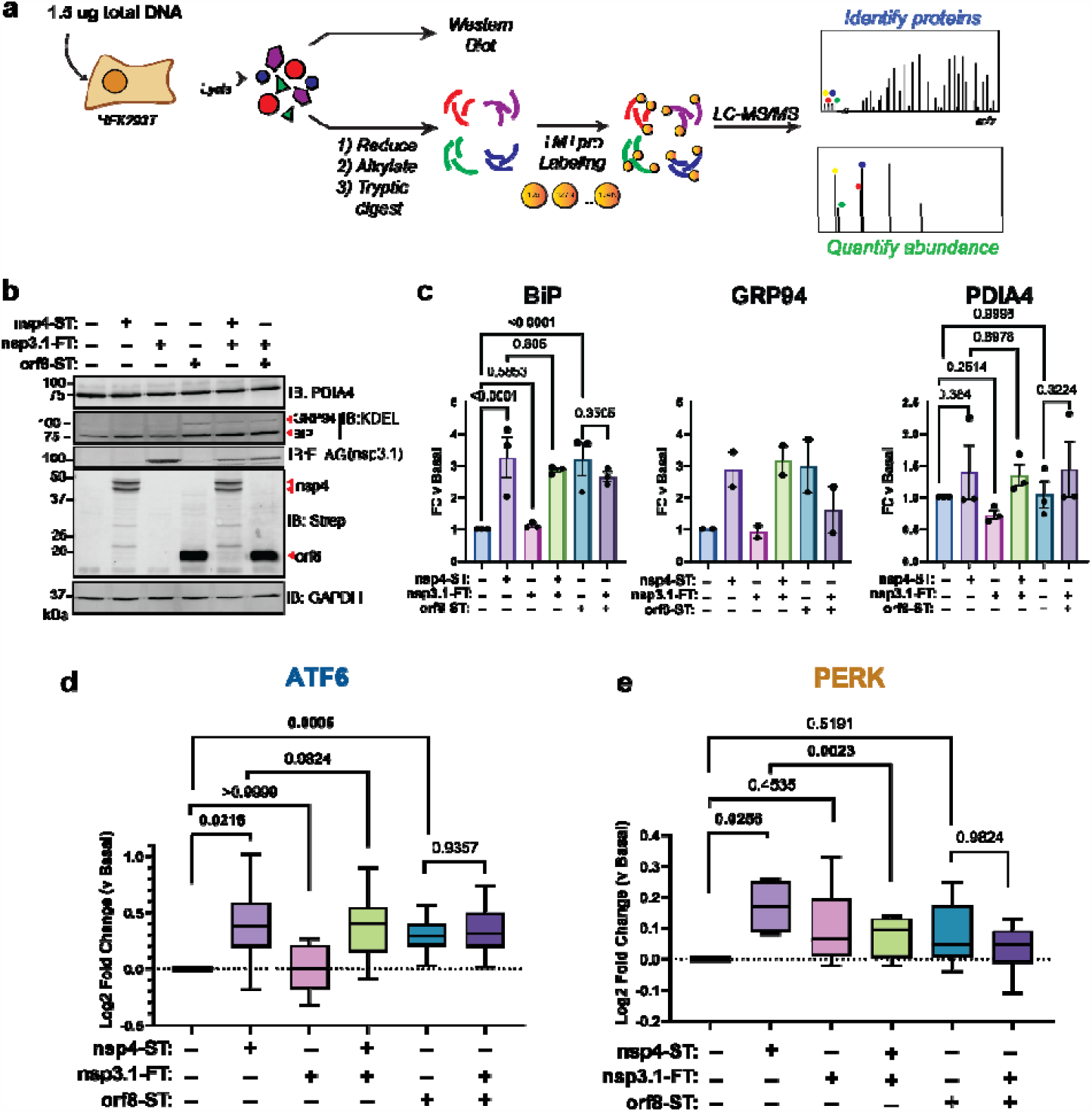
SARS-CoV-2 nsp3.1 suppresses nsp4-induced PERK, but not ATF6 activation. **a)** Experimental schematic to test the effect of SARS-CoV-2 nsp3.1 on nsp4- or orf8-induced activation of the UPR in HEK293T cells by Western blot and tandem mass spectrometry (LC-MS/MS). **b)** Representative Western blot of ATF6 protein markers (PDIA4, BiP, GRP94) and viral proteins (SARS-CoV-2 nsp4-ST, nsp3.1-FT, and orf8-ST), with GAPDH as a housekeeping gene for a loading control. See **Supplemental Fig. S5** for all quantified, original blots. **c)** Quantification of Western blots in **(b)**, normalized to GAPDH band intensities. n = 2-3, mean ±SEM, One-way ANOVA with Benjamini, Krieger and Yekutieli multiple testing correction, p<0.05 considered statistically significant. **d)** Global proteomics analysis of ATF6 protein marker levels in the presence of SARS-CoV-2 nsp4-ST, nsp3.1-FT, orf8-ST, or specified combinations. Box-and-whisker plot shows median, 25^th^ and 75^th^ quartiles, and minimum and maximum values. one-way ANOVA with Geisser–Greenhouse correction and post-hoc Tukey’s multiple comparison test was used to test significance; 1 MS run, n = 2-3 biological replicates. See **Supplemental Tables S2, S3** for mass spectrometry data set. **e)** Global proteomics analysis of PERK protein marker levels, as in **(d)**.

As expected, we found that transfection with nsp4 and orf8 individually induced increased protein expression of the ATF6 markers BiP, GRP94, and PDIA4 compared to basal conditions, as measured by Western blot (**Fig. 3b,c, Supplemental Fig. S5**). However, nsp3.1 did not lead to any notable decrease in the expression of ATF6 markers when co-expressed with nsp4 or orf8.

To more comprehensively quantify UPR protein markers, we analyzed these same lysates by TMTpro-quantitative LC-MS/MS. We found that nsp3.1 does not suppress nsp4- or orf8-induced ATF6 activation (**Fig. 3d**). Interestingly, we found that co-expression of nsp3.1 with nsp4 significantly lowers PERK marker levels compared to nsp4 alone (**Fig. 3e**), while IRE1/XBP1s activation was not significantly decreased with co-expression (**Supplemental Fig. S6**). These results demonstrate that nsp3.1 is not capable of suppressing ATF6 activation induced by other viral proteins, such as nsp4 or orf8, but can suppress PERK activation.

Curiously, we also noted nsp3.1 protein levels are lower in co-transfections with nsp4 versus GFP, as seen by both Western blotting (**Fig. 3b**) and proteomics (**Supplemental Fig. S7**). Examining global changes in the proteome with nsp4 expression showed few proteins are substantially down- or up-regulated (**Supplemental Fig. S7**). Therefore, it is likely that this is a specific effect on nsp3.1 protein levels.

### Re-analysis shows time-dependent PERK activation during SARS-CoV-2 infection

Previous work has shown that the PERK-eIF2α-ATF4 signaling pathway is activated during SARS-CoV-2 infection^3^. A re-analysis of global proteomics during a SARS-CoV-2 infection time-course in Caco-2 cells^31^ shows a moderate upregulation of PERK-induced protein markers at 6 hpi that is downregulated by 24 hpi (**Fig. 4**). This time-dependent regulation of the PERK UPR pathway induction and subsequent suppression may be explained by nsps, such as nsp3.1 and nsp4. Early activation of the PERK pathway may be important to interfere with host translation through eIF2α phosphorylation, while the later deactivation would avoid ATF4/CHOP-mediated cell death typically associated with prolonged PERK activation.

**Figure 4.**
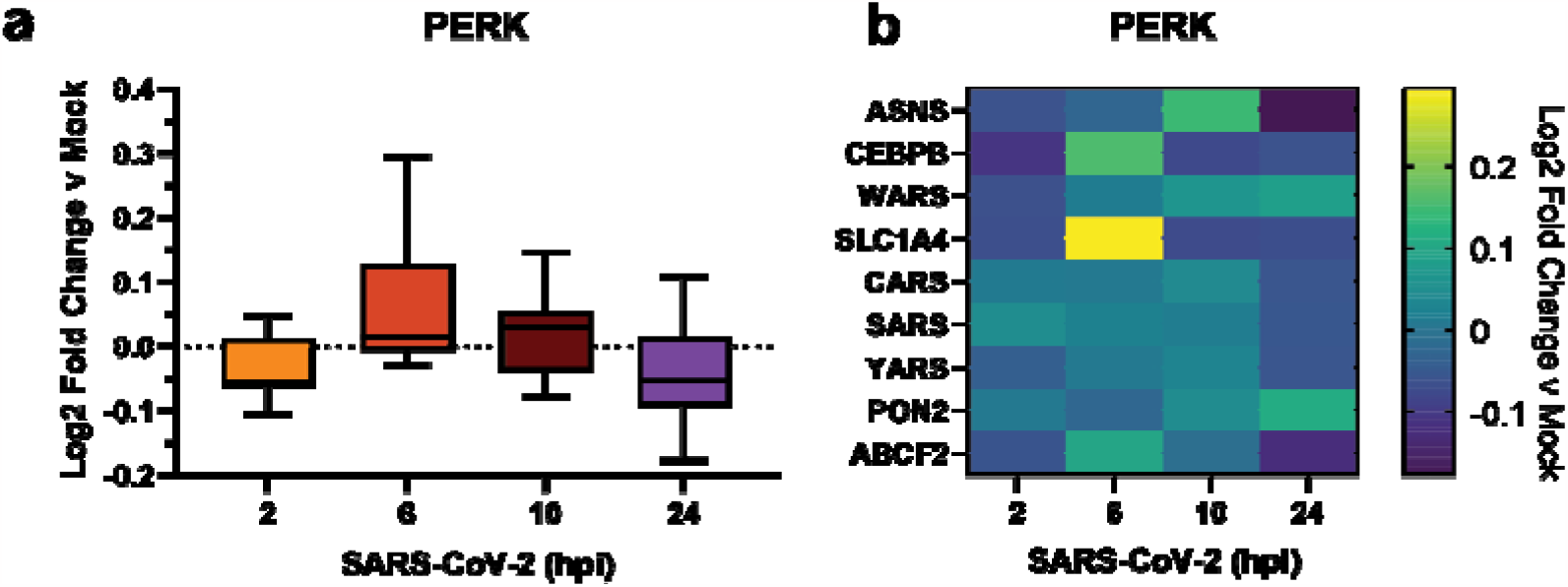
Re-analysis of PERK activation in global proteomics data of Caco-2 cells infected with SARS-CoV-2. **a)** Box-and-whisker plot of PERK protein markers during SARS-CoV-2 infection (2-24 hours post infection (hpi)) compared to mock infected samples. Box-and-whisker plot shows median, 25^th^ and 75^th^ quartiles, and minimum and maximum values. **b)** Heatmap of individual PERK protein markers as in **(a)**. Dataset published by Bojkova, et al. 2020^31^.

## Conclusions

Previous studies show that SARS-CoV-2, SARS-CoV, and MHV activate branches of the UPR to varying degrees^2–4,14,15^, however the role of nsps in this process has remained largely unexplored. Of particular interest are the nsps involved in host membrane alteration to promote infection by DMV formation, such as nsp3 and nsp4. In this work, we find that SARS-CoV-2 nsp4 activates the ATF6 and PERK branches of the UPR in HEK292T cells (**Fig. 1,2**). This is evident by both Western blotting for ATF6 markers such as BiP and GRP94 and the PERK marker CHOP (**Fig. 1**). We further harnessed the capabilities of TMTpro-based quantitative proteomics to measure pathway changes in these UPR branches. The grouped analysis of co-regulated protein sets for the individual UPR pathways enables a comparative assessment of the degree of UPR branch activation and can even detect very small levels of induction (**Fig. 2**). We also find that nsp4 activates the ATF6 pathway in A549 cells, though more moderately compared to HEK293T cells (**Fig. 2d**), indicating cell-specific effects that may be tied to variable nsp4 protein expression in different cells, or divergent regulation mechanism of the UPR. We also acknowledge that transient transfection of individual or pairs of viral proteins cannot completely reflect the complete infection milieu, however this reductionist approach allows for characterization of specific viral protein contributions to UPR modulation, as has been conducted in prior studies^3,14,15^.

In both HEK293T and A549 cells, the overall degree of UPR activation by nsp4 is lower than with the highly-toxic, global ER stressor tunicamycin, but higher than treatment with branch specific pharmacologic activators (**Fig. 2, Supplemental Fig. S4**). This is in line with the moderate UPR activation observed in viral infection, where strong, chronic UPR activation may trigger apoptosis and would likely be counter-productive for replication. Additionally, prior work has found that modest upregulation of individual UPR branches can substantially improve quality control and secretion of disease-relevant proteins, such as in light-chain amyloidosis^23,24^. Thus, the moderate UPR activation measured here may be biologically relevant to CoV-reprogramming of host cells during infection.

It is pertinent to further study the molecular mechanisms by which nsp4 trigger UPR activation. We have previously profiled a comparative interactome of nsp4 homologs^21^ and did not identify protein interactions with the ATF6 sensor protein as seen with nsp3^22^, making it unlikely that signaling occurs through a direct interaction. As a multi-pass transmembrane protein, nsp4 may be prone to misfolding or may modulate host ER membranes, which can trigger ATF6 activation in some cases^32^. Further work will be needed to define the precise molecular mechanisms by which nsp4 activates the ATF6 and PERK branches.

We previously showed that an N-terminal fragment of SARS-CoV-2 nsp3, nsp3.1, interacts with ATF6 and can suppress pharmacological activation of the ATF6 pathway^22^. We hypothesized that nsp3 and nsp4 may act in concert to tune the UPR to increase protein folding capacity while minimizing apoptotic effects of chronic activation. We found that co-expression of nsp3.1 with nsp4 or orf8 does not repress ATF6 activation but does suppress nsp4-induced PERK activation (**Fig. 3**). Time-dependent PERK activation was evident from global proteomics data during SARS-CoV-2 infection^31^ showing an increase in target protein levels during the first 6 hpi, followed by a later decline (**Fig. 4**). This tight temporal regulation suggests a role for nsp3 and nsp4 in moderating PERK signaling to permit the early signaling events (eIF2α-mediated host translational attenuation) that are beneficial for viral propagation while preventing ATF4/CHOP-mediated induction of apoptosis.

The precise mechanism by which nsp3.1 suppresses nsp4-induced PERK activation will require further investigation. SARS-CoV-2 nsp3.1 directly interacts with ATF3 ^22^, a PERK protein marker which is upregulated by ATF4 and promotes pro-apoptotic signaling^33^. This protein interaction may represent one avenue by which the PERK pathway is tuned by nsp3.1 to limit apoptosis in infected cells.

The absence of ATF6 suppression in co-expression is surprising, given that we previously showed nsp3.1 could suppress ATF6 activation by tunicamycin treatment, which potently activates the global UPR through inhibition of protein glycosylation. This may suggest that the mechanisms by which nsp4 activates the ATF6 pathway are distinct from a general tunicamycin stressor and can overcome the opposing effects of nsp3.1. Alternatively, nsp4-induced ATF6 activation may be too moderate for nsp3.1 to have any measurable effect (**Supplemental Fig. S4**). In the context of viral infection, there are multiple points of UPR induction via nsp4, orf8, Spike, ER membrane perturbation, etc. which in combination may require partial suppression by nsp3.1 to tune the host UPR and prevent host cell apoptosis.

Relatedly, co-expression of nsp4 with nsp3.1 leads to a noticeable decrease of nsp3.1 levels compared to control co-expression of GFP with nsp3.1 (**Fig. 3b, Supplemental Fig. S7**). This drop in protein levels may in part explain the lack of ATF6 suppression by nsp3.1. It is unclear how nsp4 affects nsp3.1 levels. While ATF6 activation increases production of ERAD factors to clear misfolded proteins, nsp3.1 is a cytosolic fragment of nsp3 and should not be subject to increased ERAD.

Additional questions remain regarding nsps and the UPR, such as how conserved is UPR activation across nsp4 homologs? Does nsp6, another key protein in DMV formation^16^, have a similar effect on the UPR? And how might nsp3, nsp4, and nsp6 affect the UPR in concert? Using quantitative proteomics to measure changes in UPR-induced protein expression, as we have done here, should prove a powerful and comprehensive tool in answering these questions. Lastly, previous work has shown that UPR inhibition can attenuate SARS-CoV-2^3^ and MERS-CoV^34^ infection. Viral families beyond coronaviruses, such as flaviviruses, have been shown to rely on the UPR and can be inhibited using UPR modulators^35–37^. These opposing phenotypes, in which either UPR activation or inhibition can disrupt infection, highlight the important and diverse functions the UPR plays in viral replication. Continued efforts to delineate the roles of individual viral proteins in modifying the UPR will be critical to the further development of UPR-targeting anti-virals. Our work contributes to this goal by identifying a new role for SARS-CoV-2 nsp4 as a potent UPR activator and the coordination of nsp3.1 with nsp4 to tune the PERK pathway.

## Supporting Information

Supplemental Figures of additional results (DOC)

Supplemental Table S1 of UPR pathway protein sets used for proteomics analysis (XLSX)

Supplemental Table S2 of proteins identified by LC/MS-MS in HEK293T cells (XLSX)

Supplemental Table S3 of peptides identified by LC/MS-MS in HEK293T cells (XLSX)

Supplemental Table S4 of proteins identified by LC/MS-MS in A549 cells (XLSX)

Supplemental Table S5 of peptides identified by LC/MS-MS in A549 cells (XLSX)

## Acknowledgements

We thank Dr. Nevan Krogan (University of California, San Francisco) for the pLVX-EF1alpha-SARS-CoV-2-nsp2-2xStrepTag plasmid and Dr. Joseph Genereux (University of California, Riverside) for cell stocks. We thank members of the Plate lab for their critical reading and feedback on this manuscript. Work was funded by R35GM133552 (National Institute of General Medical Sciences) and Vanderbilt University start-up funds. J.P.D. was supported by T32GM008554 (National Institute of General Medical Sciences). R.J.H.W was supported by R01DK107944 (National Institute of Diabetes and Digestive and Kidney Diseases) and R01GM121621 (National Institute of General Medical Sciences).

## Author Contributions

J.P.D, A.S., and L.P. conceptualization; J.P.D., A.S., and L.P. formal analysis; J.P.D., A.S., L.P. methodology; J.P.D., A.S., K.R.K. investigation; R.J.H.W., L.P. supervision; J.P.D., A.S., L.P. writing – original draft. J.P.D., A.S., L.P., K.R.K., R.J.H.W. writing – review & editing.

## Additional Information

The authors declare they have no conflict of interests.

