## Supplemental Information for "SARS-CoV-2 Nonstructural Proteins 3 and 4 tune the Unfolded Protein Response"

**Supplemental Tables**

**Table S1 –** UPR pathway protein sets used for proteomics analysis
**Table S2 –** Mass spectrometry protein data of HEK293T cells
**Table S3 –** Mass spectrometry peptide data of HEK293T cells
**Table S4 –** Mass spectrometry protein data of A549 cells
**Table S5 –** Mass spectrometry peptide data of A549 cells

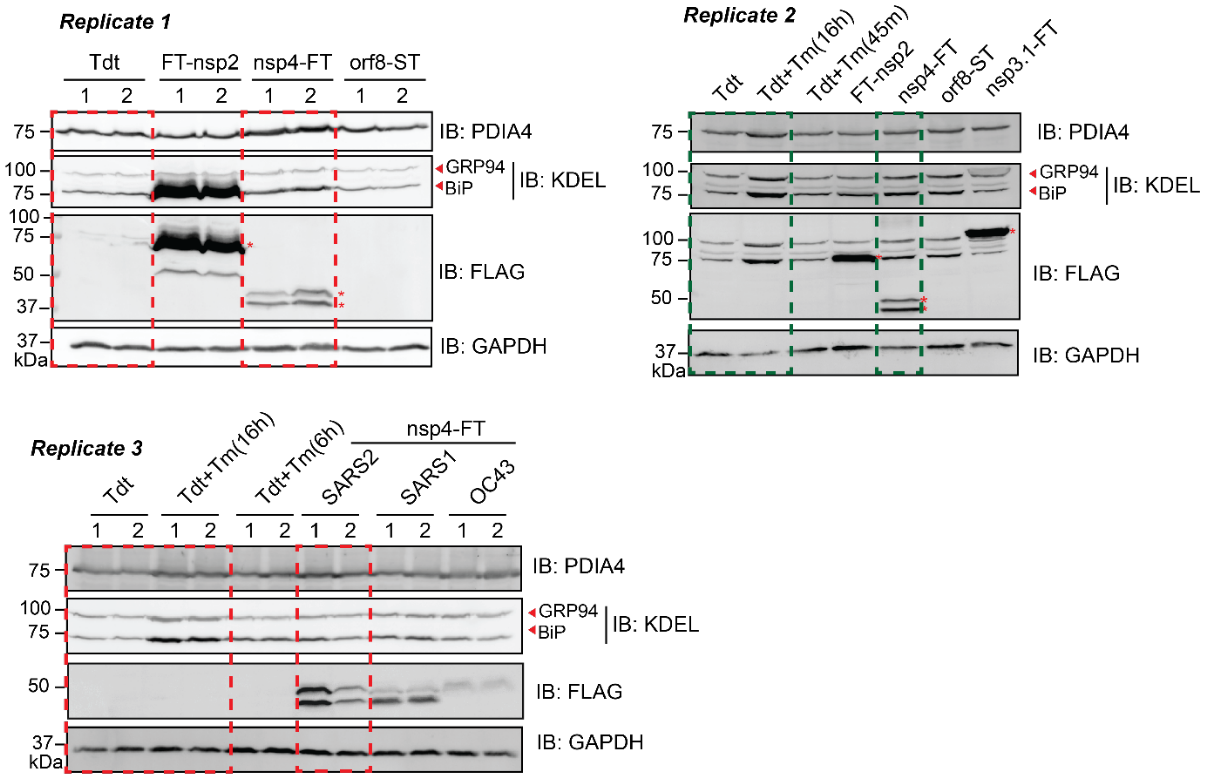


**Figure S1.** Western blot analysis of HEK293T cells transfected with specified viral proteins or Tdtomato as basal control. Tm treatment was used as a positive control. Green boxed lanes were shown for **Fig. 1c**. Green and red boxed lanes were quantified for **Fig. 1d**. Red asterisks indicate viral proteins. Tdt, tdTomato; Tm, Tunicamycin (1 µg/mL); FT, FLAG-Tag; ST, StrepTag. Corresponding full blots are shown below.

**Supplementary Fig.S1 - Replicate 1, Full blots**

IB: PDIA4, low contrast

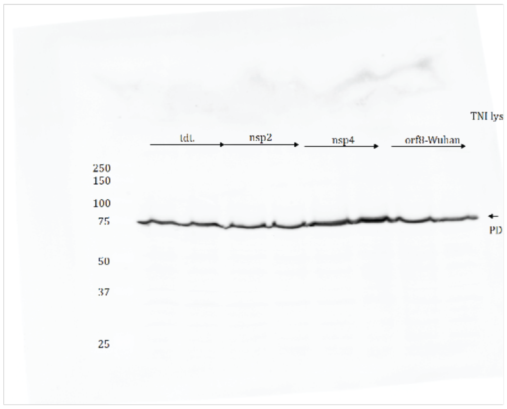


IB: PDIA4, high contrast

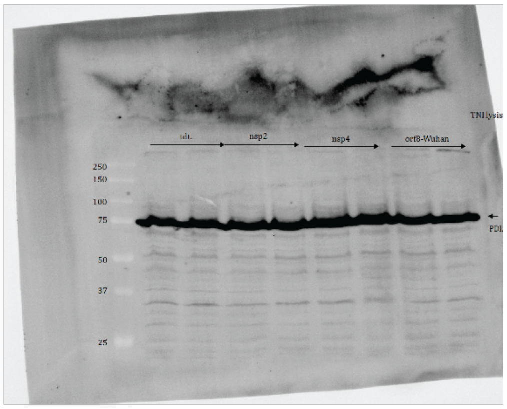


IB: FLAG

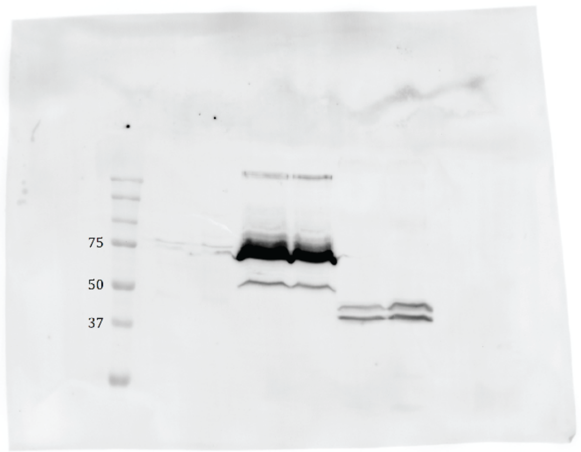


IB: KDEL

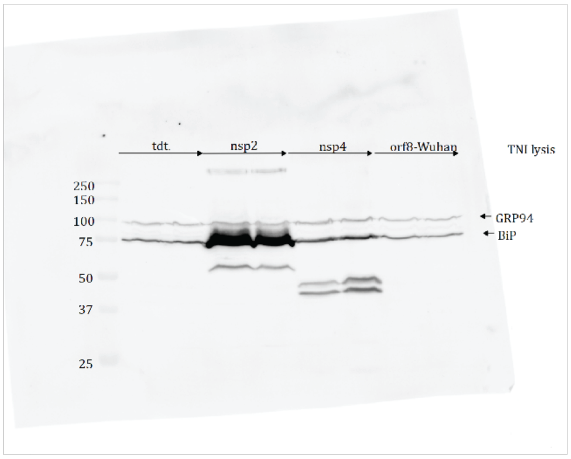


IB: GAPDH

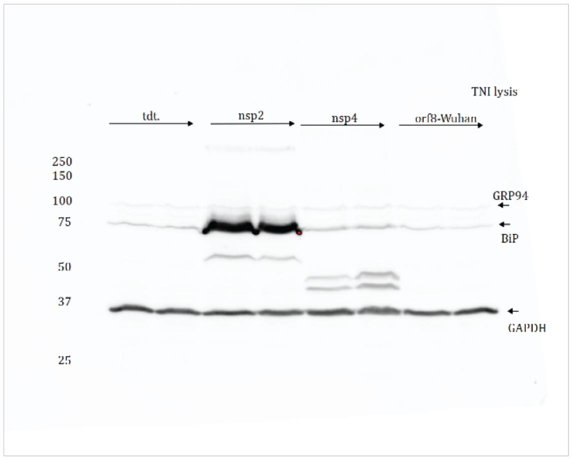

**Supplementary Fig.S1 - Replicate 2, Full blots**

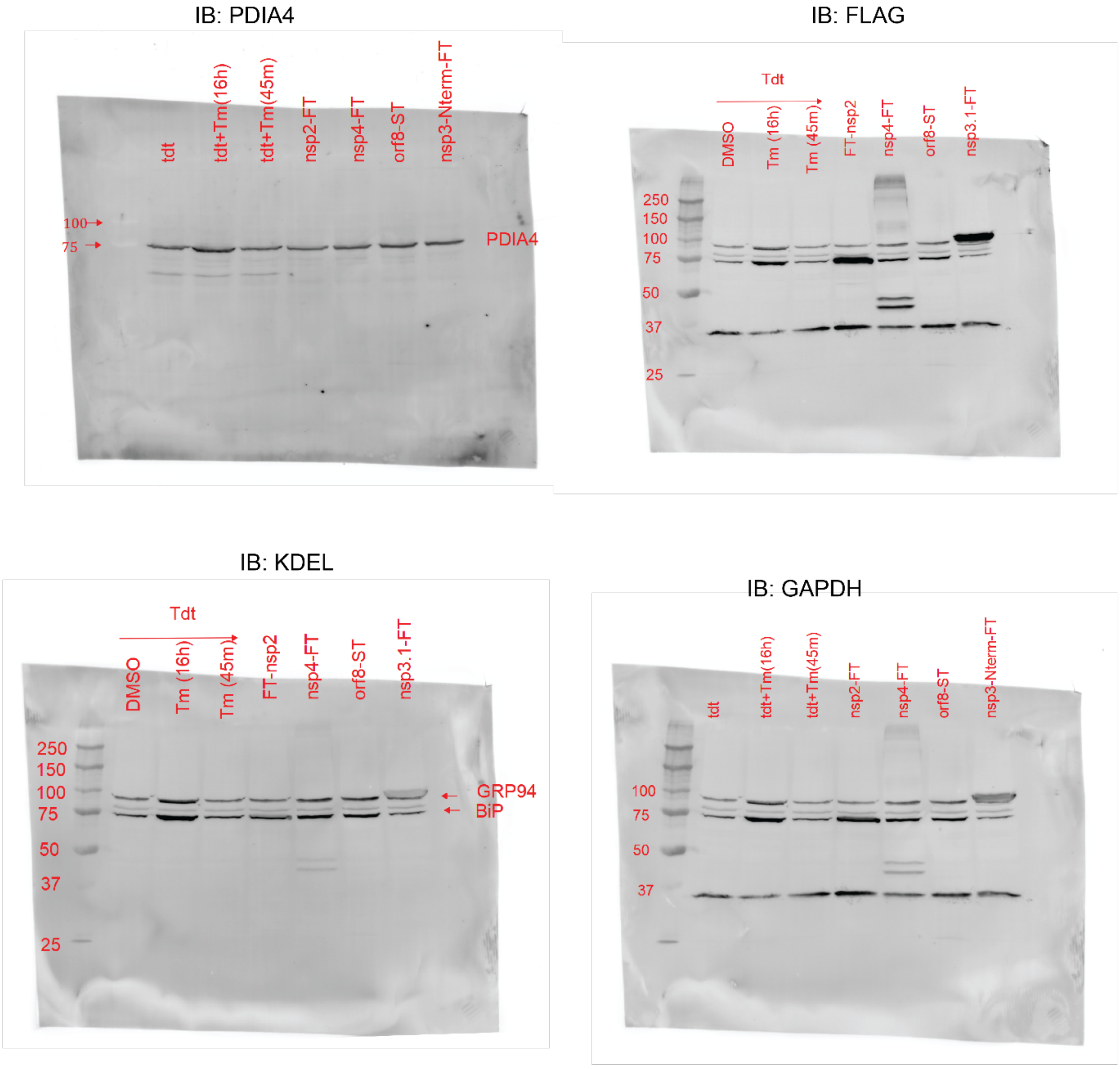

**Supplementary Fig.S1 - Replicate 3, Full blots**

IB: PDIA4

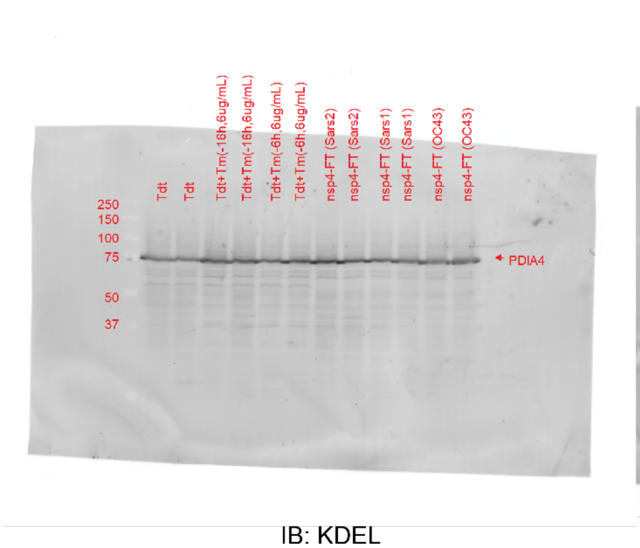


IB: FLAG

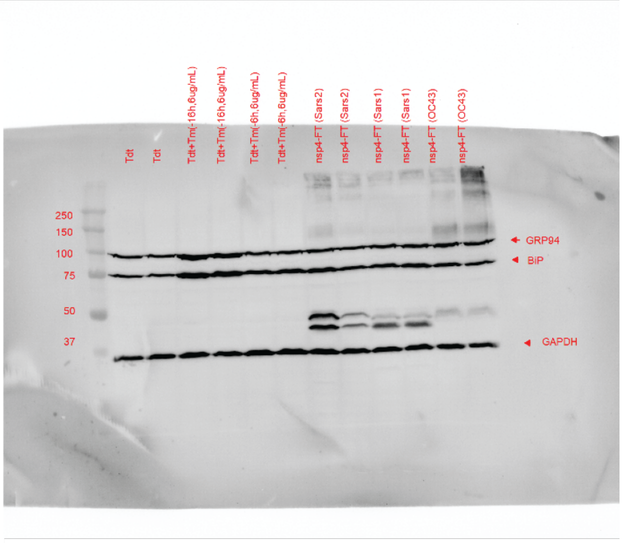


IB: KDEL

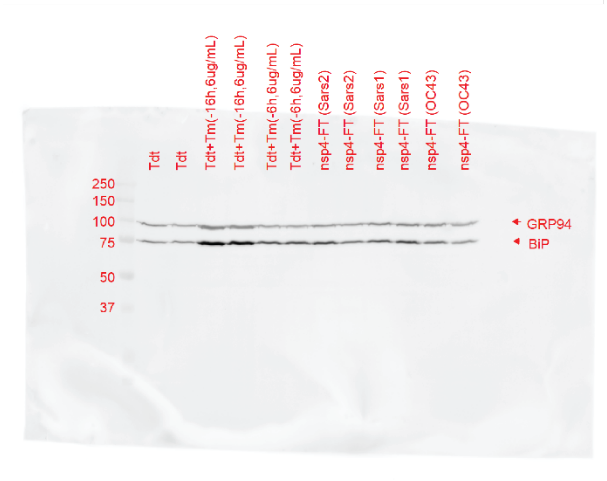


IB: GAPDH

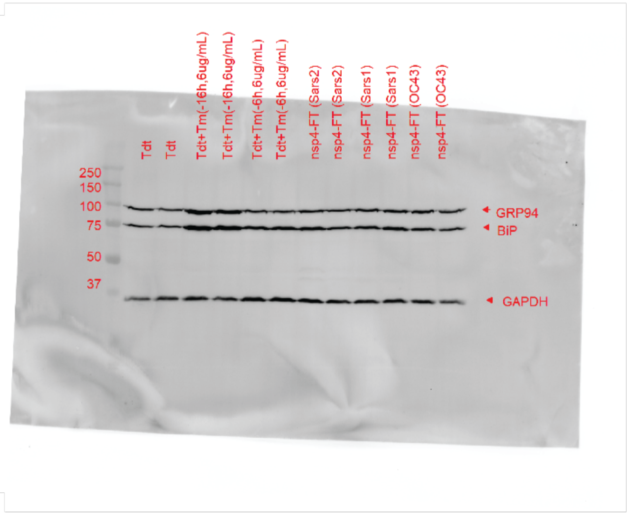

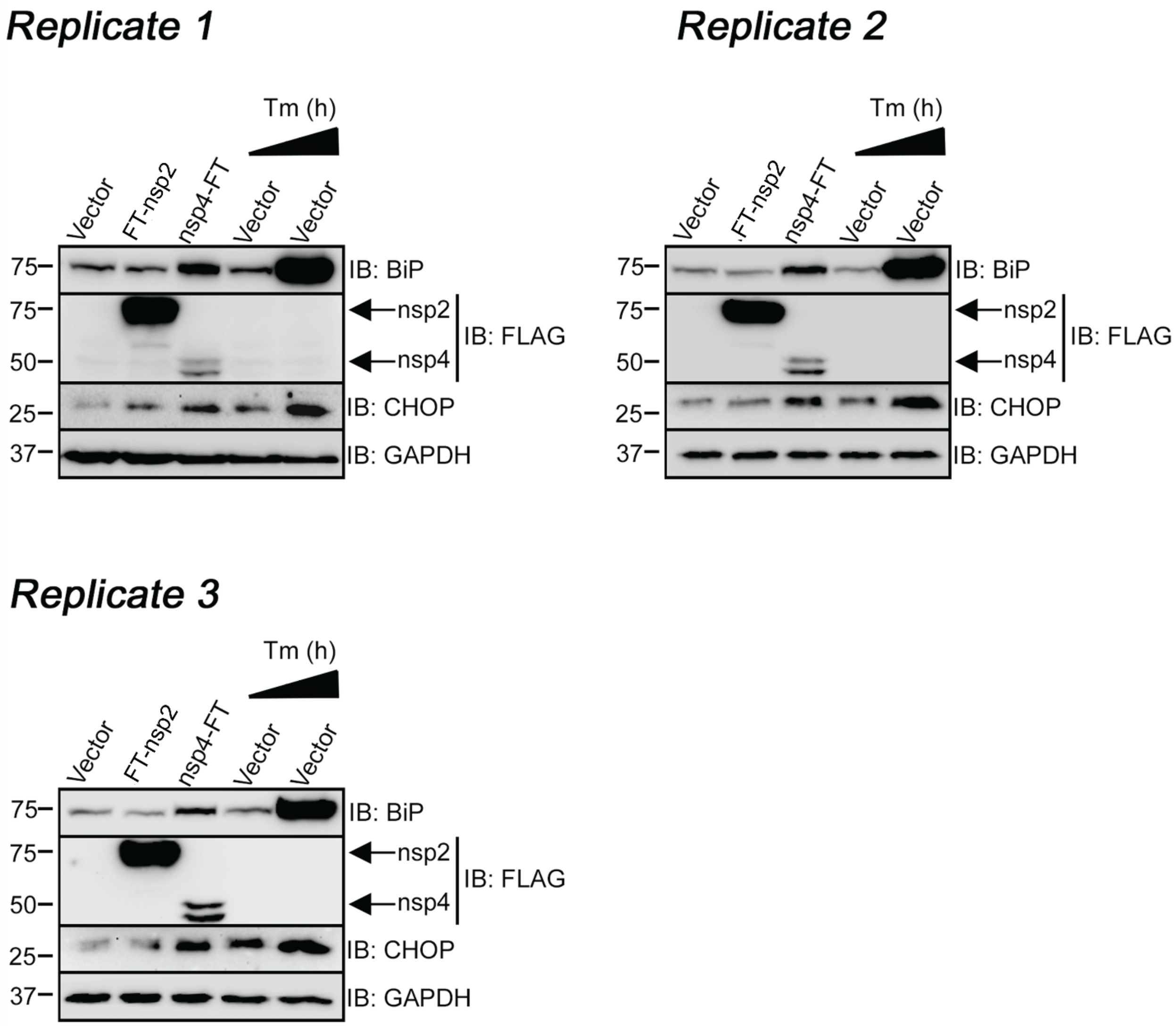


**Figure S2.** Blots quantified for **Fig. 1f**. Replicate 2 blot was shown in **Fig. 1e**. WT-HEK293T cells were transfected with empty vector or SARS-CoV-2 FT-nsp2 or nsp4-FT, with control vector samples treated for 4 or 20 h Tunicamycin (5 µg/mL). Western blotting for M2-FLAG, BiP, and CHOP shows marked upregulation of BiP and CHOP in nsp4-FT expressing cells. Corresponding full blots are shown below.

**Supplementary Fig. S2, Replicate 1 Full blots**

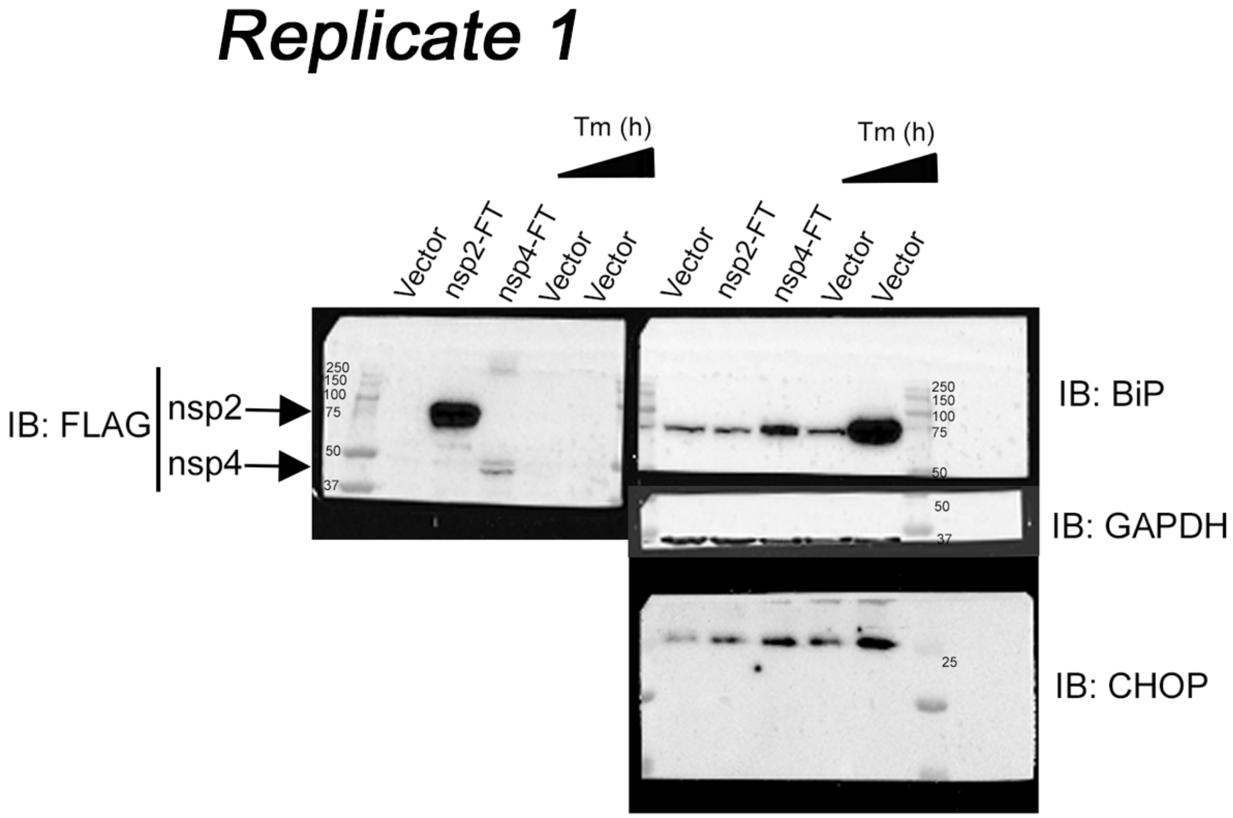

**Supplementary Fig. S2, Replicate 2 Full blots**

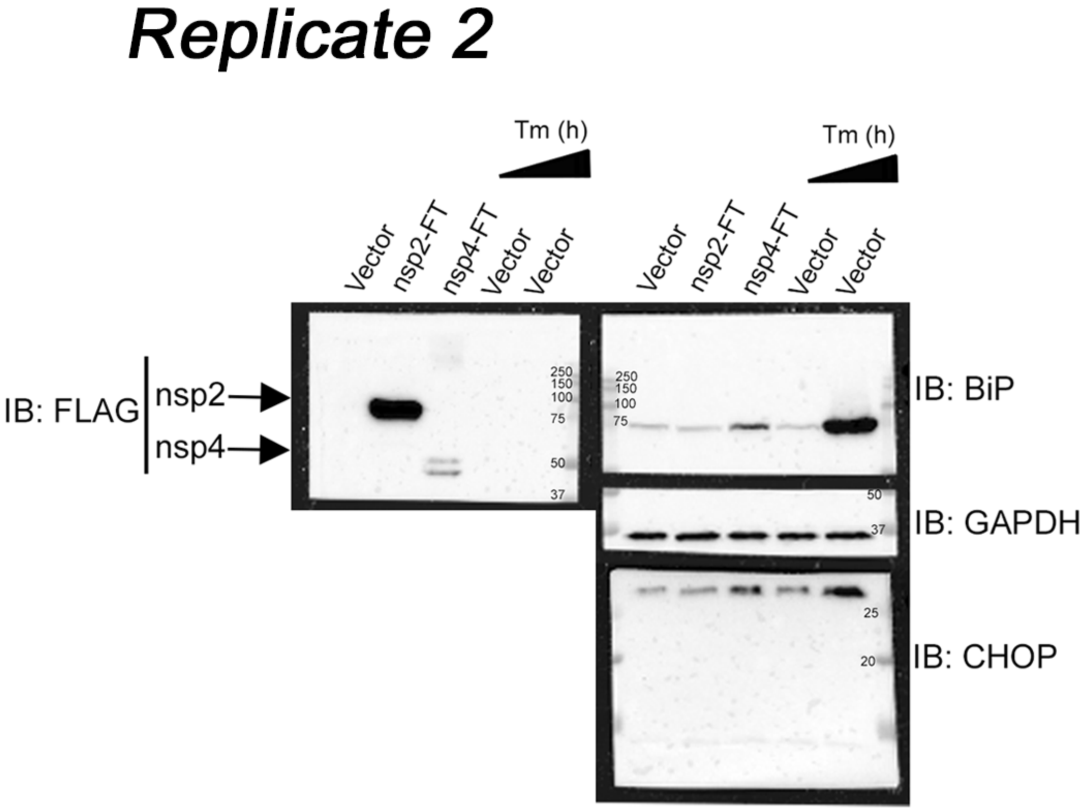

**Supplementary Fig. S2, Replicate 3 Full blots**

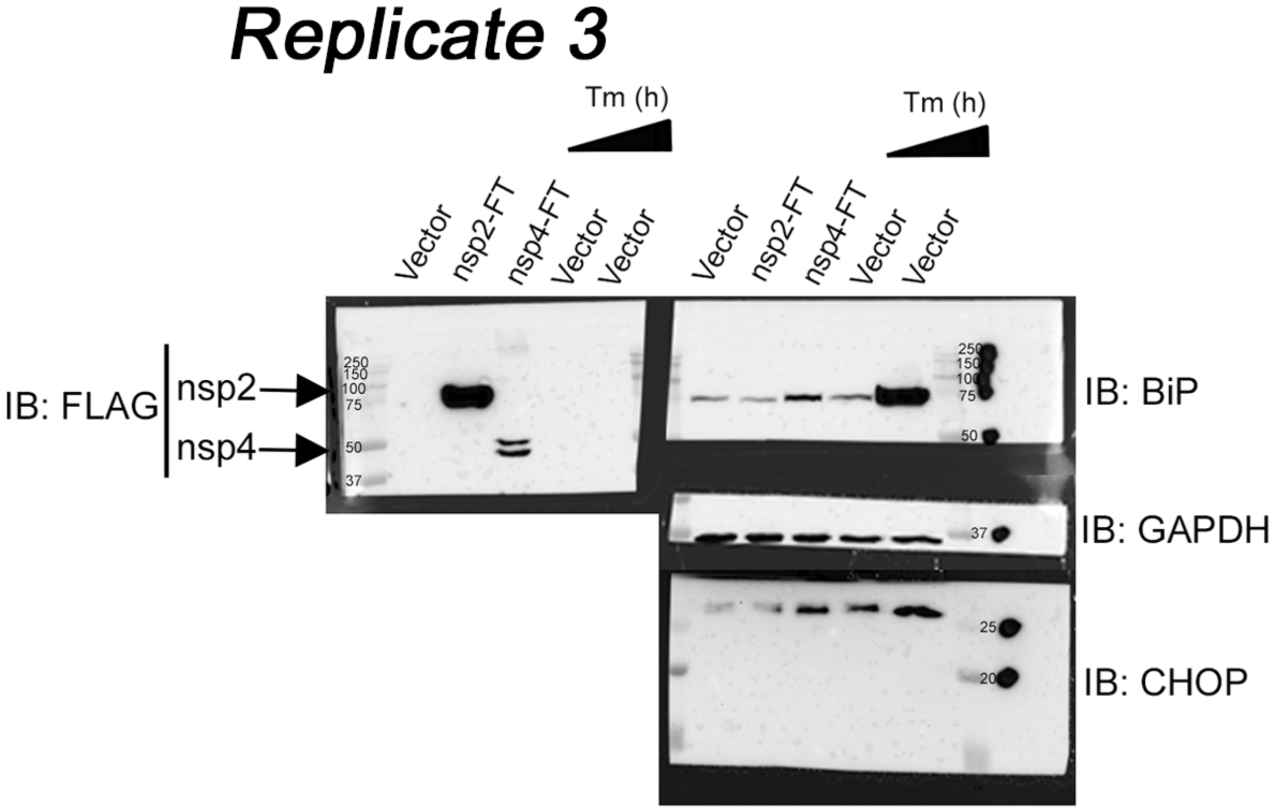

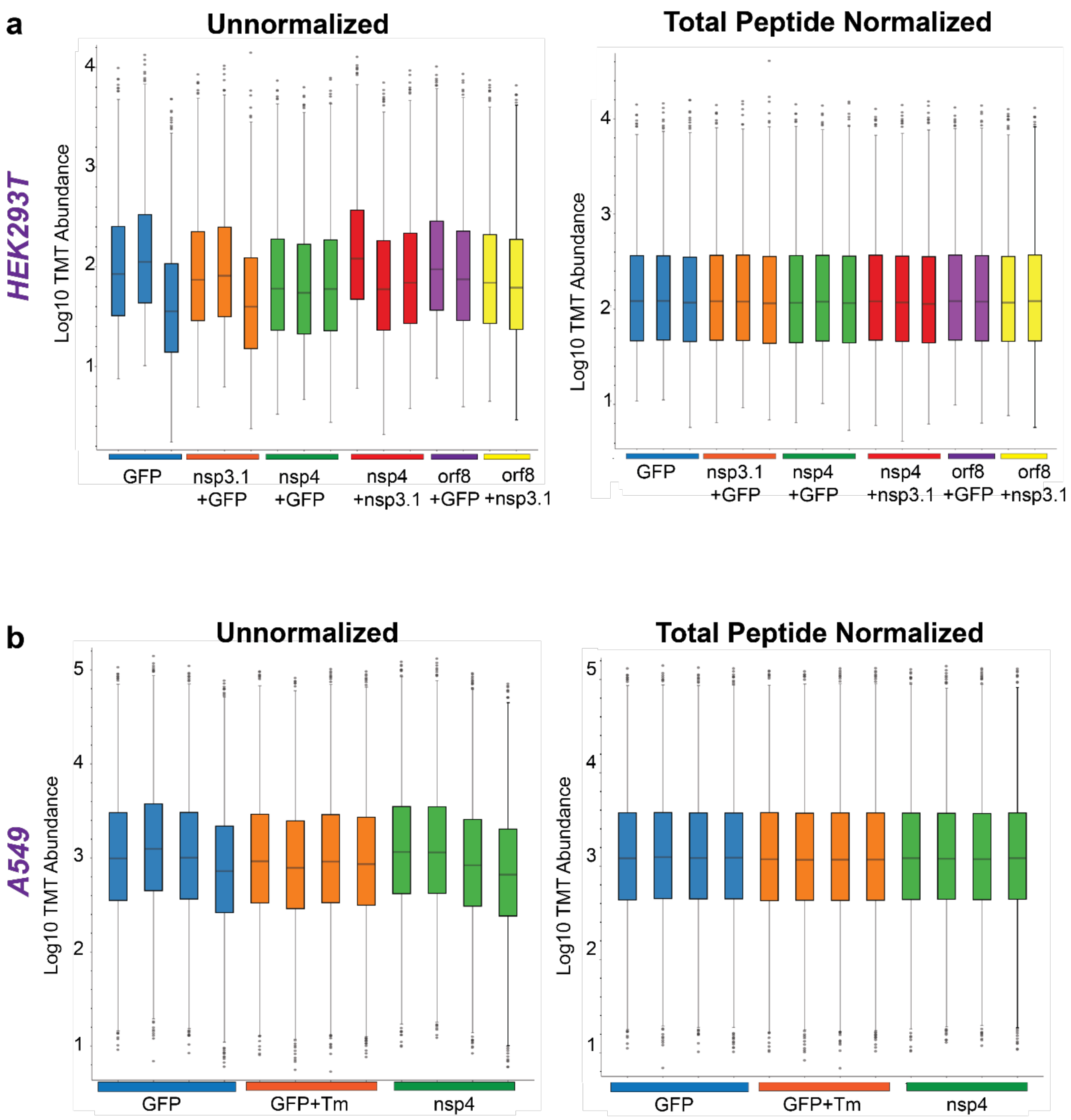


**Figure S3.** HEK293T (a) or A549 (b) cells were transfected with GFP (basal control) or corresponding viral proteins, lysates harvested and lysed at 40 h post-transfection, labeled with TMTpro isobaric labels, and analyzed by tandem mass spectrometry (LC/MS-MS). Identified proteins were normalized by total peptide amount.

**a)** Unnormalized and unnormalized log10 TMT abundances for MS experiment analyzing HEK293T UPR, corresponding to **Fig. 2,3**.

**b)** Unnormalized and unnormalized log10 TMT abundances for MS experiment analyzing A549 UPR, corresponding to **Fig. 2**.

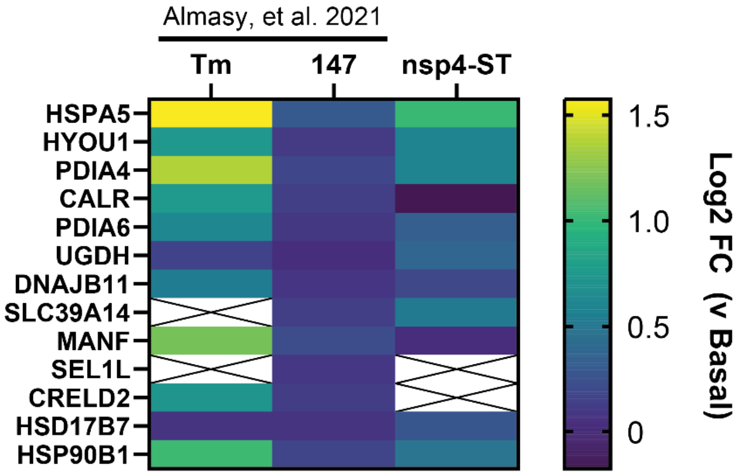


**Figure S4.** ATF6 marker log2 fold change enrichment over basal conditions in HEK293T cells with treatment of Tm (1 µg/mL, 16 h), **147** (10 µM, 16 h), or nsp4-ST expression. Samples were quantified by TMTpro-based LC/MS-MS. Basal conditions are Tdtomato transfection with DMSO for drug treatment or GFP transfection for nsp4-ST treatment. Previously published data from Almasy, Davies, Plate *MCP* 2021 is annotated[1].

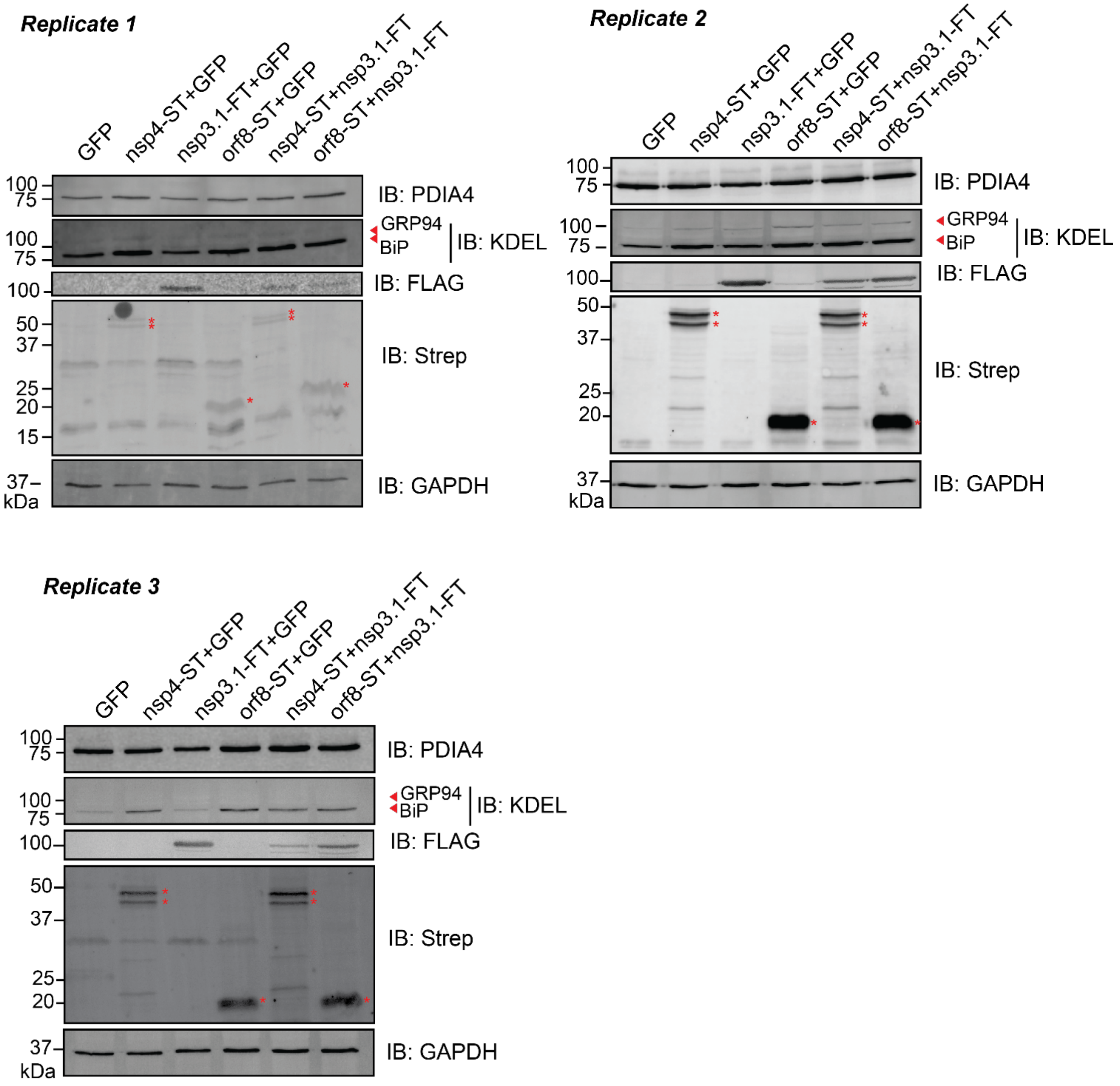


**Figure S5.** Blots quantified for **Fig. 3c**, replicate 2 was shown in **Fig. 3b**. Red asterisks indicate viral proteins. Corresponding full blots are shown below.

**Supplemental Fig.S5 - Replicate 1, Full blots**

IB: PDIA4

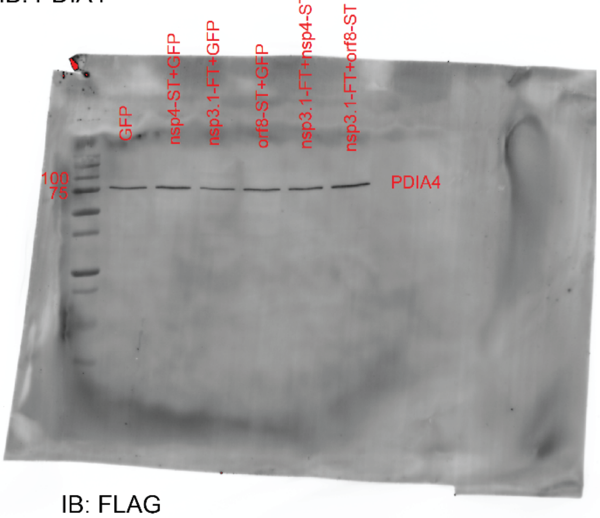


IB: KDEL

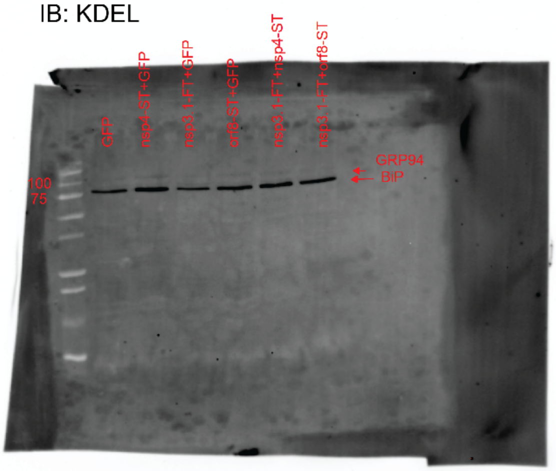


IB: FLAG

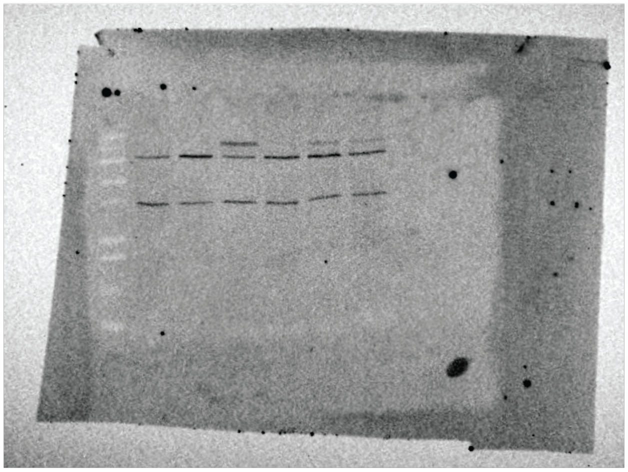

IB: Strep

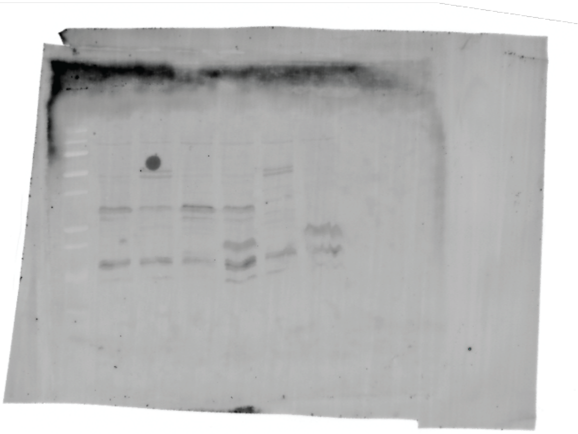

IB: GAPDH

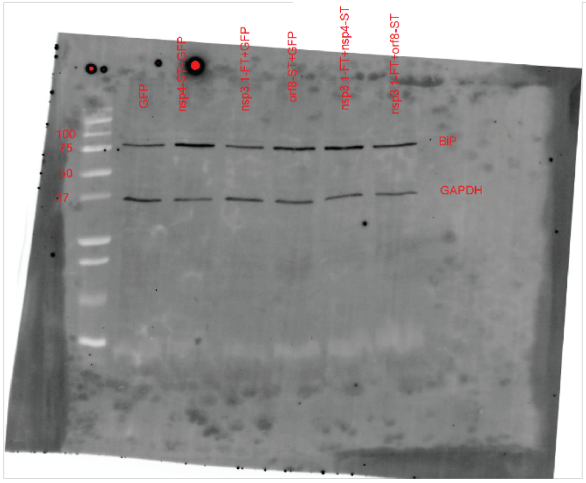

**Supplemental Fig.S5 - Replicate 2, Full blots**

IB: PDIA4

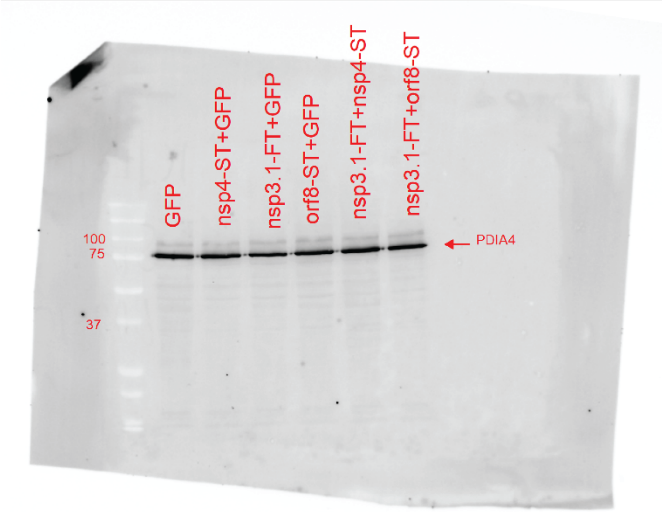


IB: KDEL

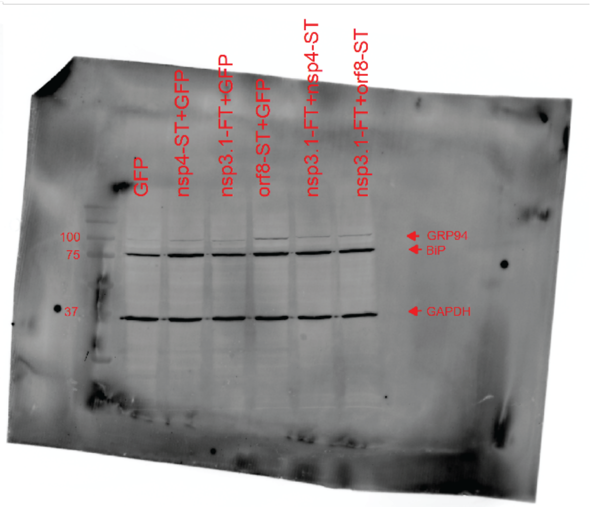


IB: FLAG

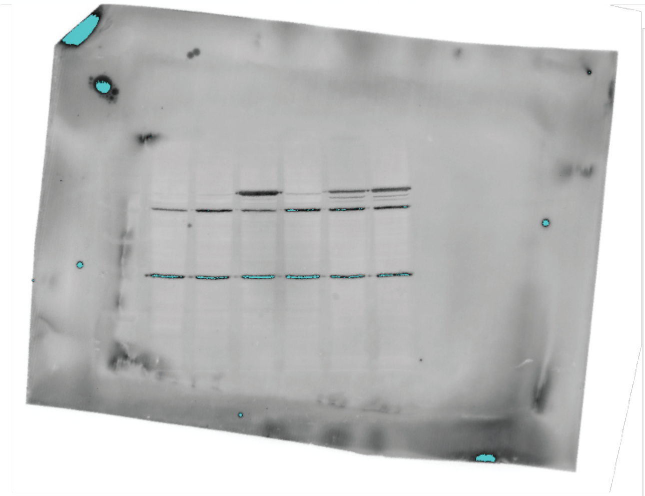

IB: Strep

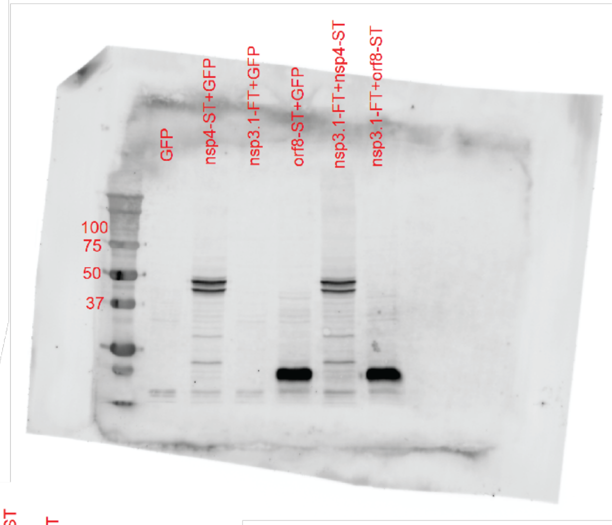


IB: GAPDH

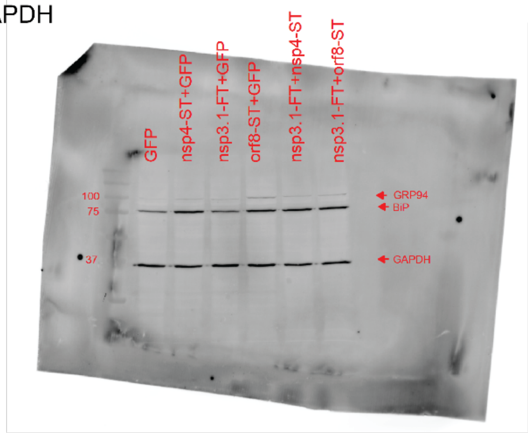

**Supplemental Fig.S5 - Replicate 3, Full blots**

IB: PDIA4

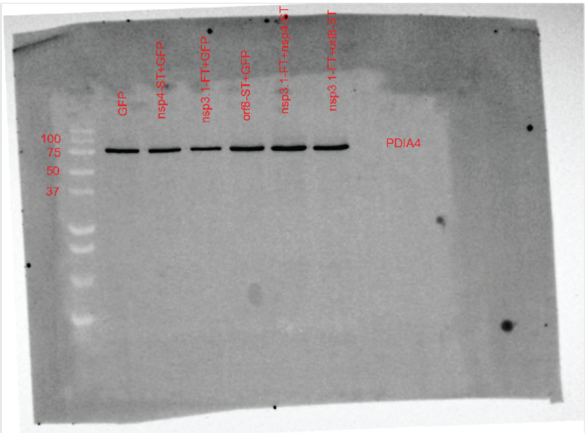


IB: KDEL

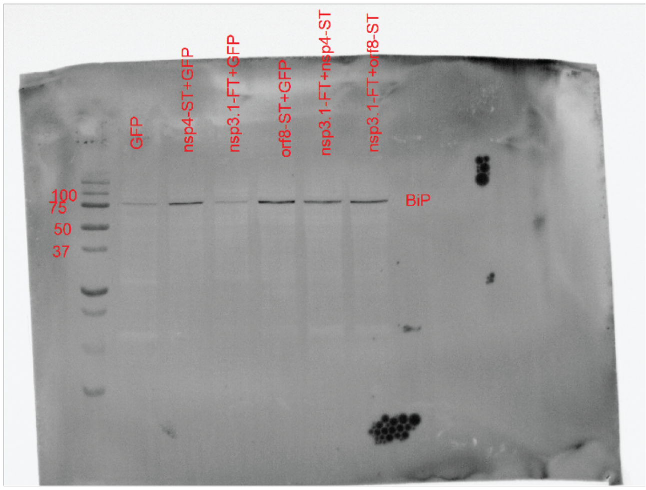


IB: FLAG

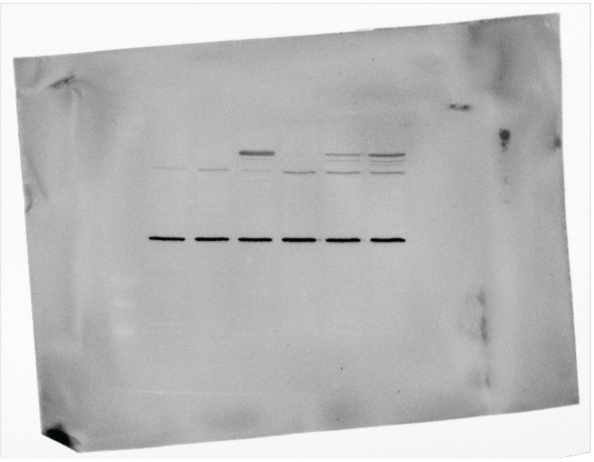

IB: Strep

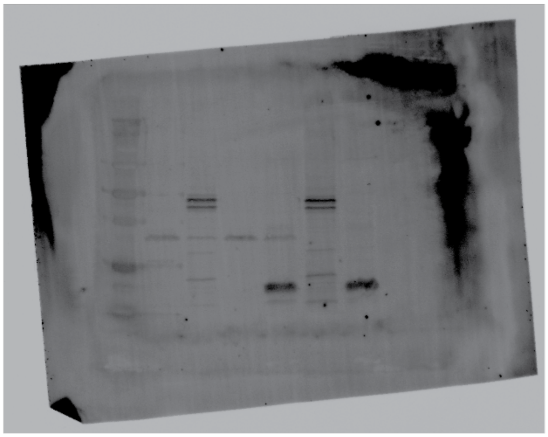

IB: GAPDH

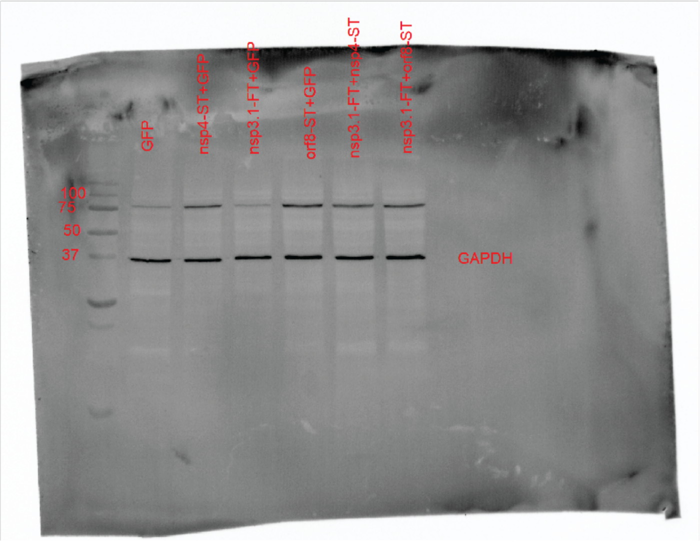

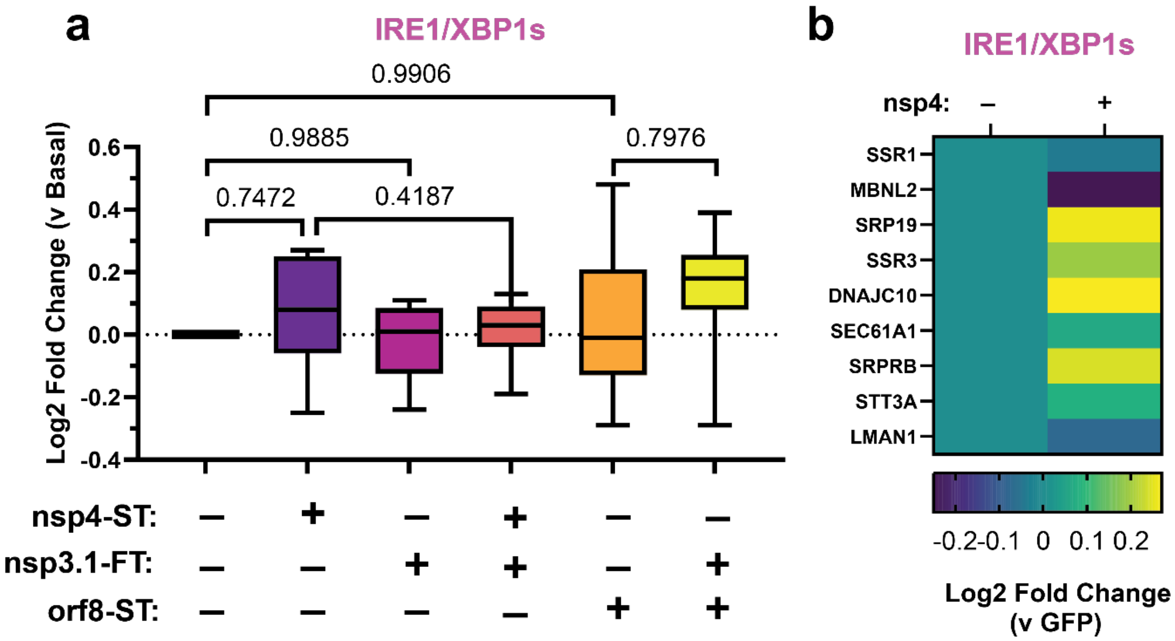


**Figure S6.**

- **a)** Global proteomics analysis of IRE1/XBP1s protein marker levels in the presence of SARS-CoV-2 nsp4-ST, nsp3.1-FT, orf8-ST, or specified combinations. Box-and-whisker plot shows median, $25^{th}$ and $75^{th}$ quartiles, and minimum and maximum values. one-way ANOVA with Geisser–Greenhouse correction and post-hoc Tukey's multiple comparison test was used to determine significance, $p<0.05$ considered significant; 1 MS run, n = 2-3 biological replicates. See **Supplemental Tables S2, S3** for mass spectrometry data set.
- **b)** Individual IRE1/XBP1s pathway protein markers in the absence or presence of nsp4-ST, from **(a)**.

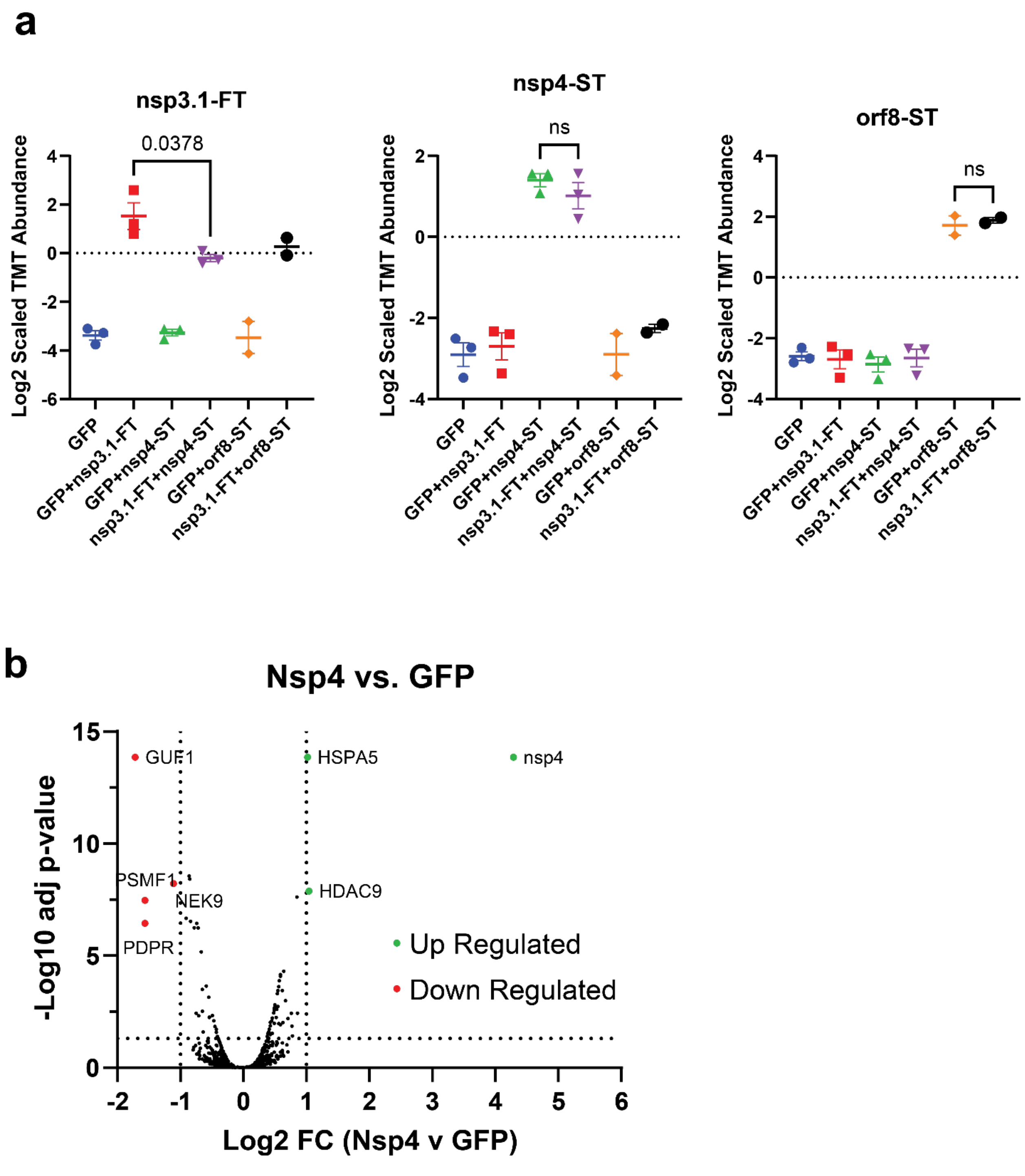


**Figure S7.**

**a)** SARS-CoV-2 nsp3.1-FT, nsp4-ST, and orf8-ST protein abundances (log2 scaled TMT abundances) in experiment corresponding to **Fig. 3.** n = 2-3 biological replicates, 1 MS run, T-test for significance between indicated samples, *p*-value annotated.

**b)** Volcano plot of global proteome in HEK293T cells in the presence or absence of SARS-CoV-2 nsp4-ST (corresponding to **Fig. 2,3**). Cut-offs indicate Log2 FC < -1 or > 1, p-value < 0.05. T-test was used for significance testing, with testing for multiple corrections.
